# Reprogramming *Yarrowia lipolytica* metabolism for efficient synthesis of itaconic acid from flask to semi-pilot scale

**DOI:** 10.1101/2023.07.17.549194

**Authors:** Jing Fu, Simone Zaghen, Hongzhong Lu, Oliver Konzock, Naghmeh Poorinmohammad, Alexander Kornberg, Deni Koseto, Alexander Wentzel, Francesca Di Bartolomeo, Eduard J Kerkhoven

## Abstract

Itaconic acid is an emerging platform chemical with extensive applications. It is currently produced by *Aspergillus terreus* through biological fermentation. However, *A. terreus* is a fungal pathogen and needs additional morphology controls, and therefore the production remains problematic. Here, we reprogrammed the GRAS yeast *Yarrowia lipolytica* metabolism for competitive itaconic acid production. After redirecting the flux of lipid accumulation as carbon sink, we evaluated itaconic acid production both inside and outside the mitochondria, and fine modulated its synthetic pathway. We then mimicked the regulation of nitrogen limitation in nitrogen replete conditions through down regulation of IDH by weak promoter changing, RNAi, or CRISPRi. Ultimately, we optimized fermentation parameters for fed-batch cultivations, and produced itaconic acid with titres of 130.1 g/L in 1L bioreactors and 94.8 g/L in a 50L bioreactor on semi-pilot scale. Our finds provide effective approaches for harnessing GRAS microorganism for competitive industrial itaconic acid production.

## Introduction

Itaconic acid (IA) is a promising platform chemical with extensive applications, and it is included in the top 12 building block chemicals ^1^ that can be converted into varied valued bio-based products. Through crosslinking, IA and its derivatives supported the synthesis of many innovative polymers with remarkable properties ^2, 3^, e.g. shape memory polymers ^4^, polymeric hydrogels used for targeted drug delivery ^5, 6^, and material against bacterial infections ^7, 8^. In addition, IA could be secreted by mammalian immune cells as part of the immune response ^9, 10^. Recently, IA was found acting as a signalling molecule in the immunomodulation ^11^, and it was a crucial anti-inflammatory metabolite to limit inflammation and modulate type I interferons ^12^, exhibiting potential applications in the biomedical field ^13^.

Current commercial itaconic acid production prefers biological fermentation over chemical synthesis ^14^, while the microorganisms used are not ideal for economic IA production at industrial level. The production is primarily through the use of the filamentous fungus *Aspergillus terreus*, which has been reported to produce 160 g/L IA ^15^. However, its IA production remains challenging. First, *A. terreus* is a fungal pathogen that can cause lethal infections ^16^. Se cond, morphological control has a significant effect on the IA production ^17^ and maintaining the required growth-form pellet needs careful setting of fermentation parameters ^18^. IA production has additionally been reported to be severely inhibited when the concentration of Mn^2+^ exceeded an extremely low level of 3 μg/L ^15, 19^, while a cation exchange treatment of the analytical grade glucose ^19^ or carefully addition of calcium ^20, 21^ or copper ions ^22^ to antagonize the effect of manganese ions was necessary to afford highest yield. The pathogen status and additional controls could increase operational cost and risks of failed batches ^23^. Meanwhile, another promising microorganism *Ustilago maydis*, as a ubiquitous pathogen of corn ^24^, has been reported to produce more than 220 g/L itaconate as solid calcium salt form ^23^. However, that fermentation requires manual addition of solid calcium carbonate as liquid suspension or powder whenever the pH dropped below 6.2. Meanwhile, calcium itaconate as the final product requires in-situ precipitation to alleviate product inhibition, placing significant hurdles for scale up. Besides these two native IA producers, numerous heterologous IA producers have been engineered, such as *Escherichia coli* ^25, 26, 27^, *Saccharomyces cerevisiae* ^28^, *Corynebacterium glutamicum* ^29^, *Aspergillus niger* ^30, 31^ and *Yarrowia lipolytica* ^32, 33, 34^. However, their IA production has not been satisfactory, and production parameters have been too far from commercialization ^35^.

*Y. lipolytica* is recognized as a “recommended biological agent for production purposes” by the European Food Safety Authority^36^, and has been granted Generally Recognized As Safe (GRAS) ^35^ status to various commercial scale processes performed with its involvement ^37^. It has been gaining traction as a biotechnologically relevant cell factory and is arguably regarded as the most promising non-conventional yeast biocatalyst ^37^. The main advantage to employ *Y. lipolytica* as IA producer, other than *S. cerevisiae* with more genetic manipulation tools, is due to its considerable metabolic flux to citric acid (CA)/isocitrate acid (ICA) that can be redirected to IA via the intermediate cis-aconitate. Significantly, its pseudo-hyphae formation can be completely abolished by deleting mhy1 ^38^, which is superior compared to *A. terreus* requiring special morphological control ^17^. *Y. lipolytica* can synthesize and accumulate large quantities of CA (>100 g/L) and its derived lipids^39^ when triggered by nutrient restriction ^40, 41^, such as nitrogen limitation (NL), phosphate limitation (PL), or sulfur limitation (SL). As an overflow metabolite, IA production is mainly synthesized during the non-growth ^13, 18^ phase in *A. terreus* and *U. maydis* with abundant carbon source present, and a serious shortage of phosphate or nitrogen as a fundamental prerequisite ^13^. However, in *Y. lipolytica*, there have been conflicting reports about whether IA production is growth associated or not ^32, 33, 34^.

In this study, we engineere *Y. lipolytica* for efficient IA production by reprogramming its metabolism. We establishe a competitive IA producer and tested it from shake flask to semi-pilot scale bioreactor, addressing the following aspects within 4 modules. In Module 1, we achieve enhanced cis-aconitate supply by redirecting the instinctive carbon flux away from the competing lipid storage, and by interrupting the glyoxylate cycle and NADP^+^ dependent isocitrate dehydrogenase (IDP). In Module 2, we introduce the IA synthetic pathway and enhance it by compartmentalization strategy and fine-tuned expression levels of key enzymes. In Module 3, we successfully decouple high flux of cis-aconitate and NL condition, by mimic NL regulations without NL.

The optimized carbon distribution between cell growth and product in nitrogen replete conditions is achieved by down regulating expression of NAD+ dependent isocitrate dehydrogenase (IDH). Ultimately, we optimised fermentation parameters in Module 4, and we produced 130.1 g/L of IA in 1L bioreactors and 94.8 g/L of IA in a 50L bioreactor with fed-batch. Overall, our work presents a significant leap and provide effective approaches towards harnessing *Y*.*lipolytica* for competitive biotechnological production of itaconic acid.

## Results

### Reduce flux in sink pathways to enhance cis-aconitate supply

Ahead of establishing heterologous itaconic acid (IA) biosynthesis, a reduction in the fluxes of alternative citric acid (CA) and isocitric acid (ICA) sink pathways should enhance the accumulation of the IA precursor cis-aconitate.

To block CA-derived lipid accumulation, three acyltransferase coding genes, *DGA1, DGA2*, and *LRO1* involved in TAG accumulation, and the Are1p coding gene *ARE1* involved in sterol esterification, were deleted in the starting strain OKYL029, a strain with mhy1 deletion to prevent pseudo-hyphae formation (Fig. 2a). Under nitrogen limitation (NL), when lipid accumulation would customarily peak, the resulting JFYL007 strain strongly decreased lipid accumulation per CDW while displaying a decreased μ_max_ (Fig. 2b). No lipid droplets were observed by microscopy (Fig. 2c). Levels of all five major fatty acids decreased significantly (Fig. 2d). As we later found that nitrogen replete (NR) conditions afforded better IA productivity, and *Y. lipolytica* ordinarily still stores low amounts of lipids at NR conditions ^42^, JFYL007 was cultivated in NR medium and cis-aconitate accumulation indeed increased (Fig. 2e), indicating that the lipid accumulation flux was redirected to cis-aconitate.

To reduce flux in the ICA sink pathways, ICA utilization by the glyoxylate cycle and isocitrate dehydrogenation were disrupted. For isocitrate lyase *ICL1* and *ICL2* were deleted to increase cis-aconitate accumulation (Fig. 2e). Meanwhile, disruption of isocitrate dehydrogenation involved two isoenzymes, NAPD^+^-dependent IDP and NAD^+^-dependent IDH. The deletion of *IDP* only modestly increased cis-aconitate accumulation, while the final biomass decreased. Co-deletion of *ICL1, ICL2* and *IDP* significantly increased cis-aconitate accumulation. Contrastingly, five attempts to delete the open reading frames of IDH subunits coding genes *IDH1* or *IDH2* failed, suggesting that the NAD^+^ dependent IDH is essential for *Y. lipolytica* survival on YPD medium plates.

Notably, in NR conditions, no detectable CA or ICA accumulated in any of the above strains. In conclusion, the increased precursor supply, and absence of overflow metabolites, rendering them as promising chassis for IA production.

### Enhance IA production by optimizing its synthetic pathways

In *Aspergillus terreus*, mitochondrial aconitase (ACO) converts CA into ICA in a 2-step reaction that yields cis-aconitate as intermediate, which is transported to the cytosol for conversion to IA by cis-aconitate decarboxylase (CAD) ^43^. Contrastingly in *Ustilago maydis*, cytosolic cis-aconitate is first converted by aconitate isomerase (ADI) to trans-aconitate, which is transformed to IA by trans-aconitate decarboxylase (TAD) ^35^. These two IA synthetic routes, both outside and inside mitochondria, were constructed and evaluated in *Y. lipolytica* (Fig. 1, module 2).

**Fig. 1.**
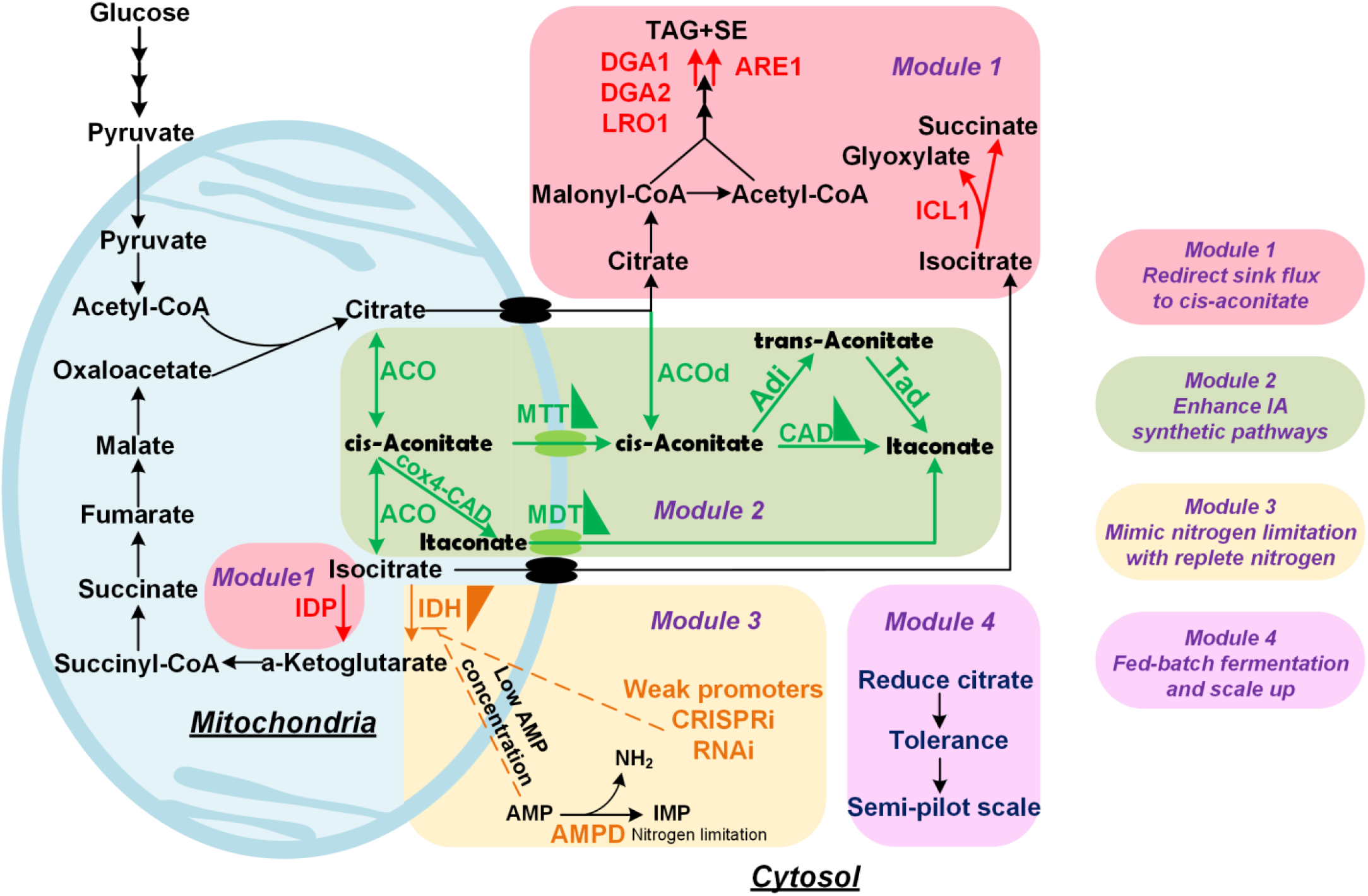
Reprogramming *Y. lipolytica* metabolism for efficient synthesis of itaconic acid from flask to semi-pilot scale. The whole work could be divided into 4 modules. Module 1, to redirect sink flux, there were mainly two nodes, citrate and isocitrate. *DGA1, DGA2, LRO1* and *ARE1* were deleted to block the TAG and SE accumulation. *ICL1* was deleted to block the glyoxylate cycle, and IDP was deleted to reduce the utilization of isocitrate. Module 2, enhance IA synthetic pathways. Module 3, the mechanism in NL was mimicked by down regulation of IDH by weak promoter exchange, CRISPRi and RNAi. Module 4, the scale-up from deep-well plates to bench top bioreactors and semi-pilot scale fermentation. ACO, aconitase; ACOd, aconitase without mitochondrial leading sequence; ADI, aconitate isomerase; AMPD, AMP deaminase; ARE1, Acyl-CoA:sterol O-acyltransferase; CAD, cis-aconitate decarboxylase; DGA1, Acyl-CoA diacylglycerol O-acyltransferase 1; DGA2, Acyl-CoA diacylglycerol O-acyltransferase 2; ICL1, isocitate lyase; IDH, NAD^+^ dependent isocitrate dehydrogenase; IDP, NADP^+^ dependent isocitrate dehydrogenase; LRO1, phospholipid:diacylglycerol acyltransferase; MDT, mitochondrial decarboxlic transporters; MTT, mitochondrial tricarboxlic transporters; TAD, trans-aconitate decarboxylase.

To test whether the earlier cis-aconitate enhancing strategy (Fig 2e) could enhance IA production, a codon optimized *A. terreus* CAD gene was introduced. Blocking lipid accumulation increased IA production from 0.027 to 0.049 g/L, while while blocking glyoxylate cycle yielded 0.126 g/L of IA (Fig. 3a). The co-deletion of ICL1/2 and IDP resulted in the highest IA accumulation of 0.184 g/L in JFYL033, which is consistent with their enhanced cis-aconitate accumulation (Fig. 2e). As several reports have shown that subcellular localization of metabolic pathways can efficiently increase product conversion ^44^, the CAD was targeted to the mitochondria by fusing it with the mitochondrial leading sequence of the Cytochrome coxidase subunit 4 (cox4). As anticipated, more IA (0.221 g/L) was produced with mitochondrial CAD, suggesting more efficient transformation to IA (Fig.3a) (Fig. 3a).

**Fig. 2.**
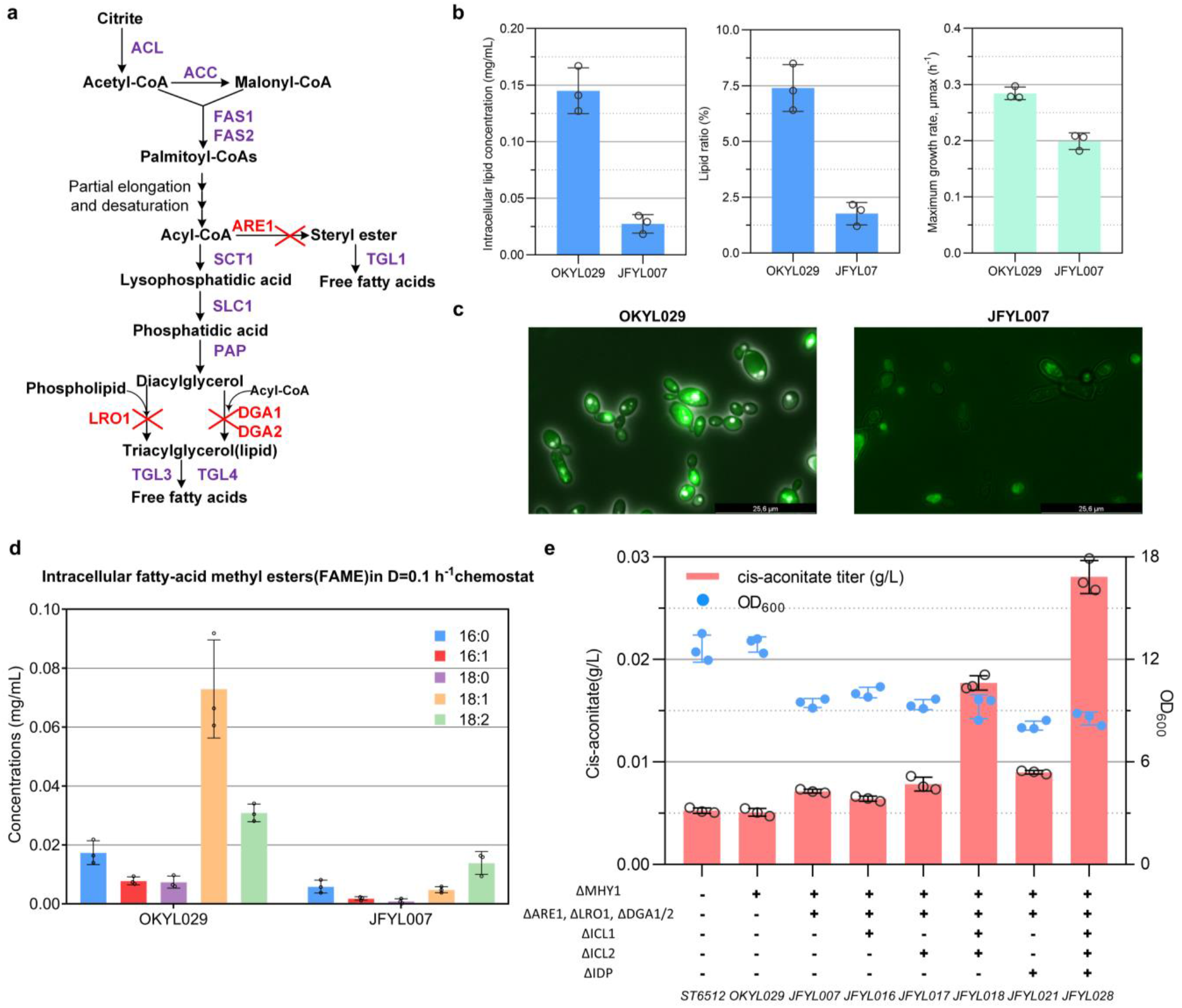
Redirect sink pathways to increase cis-aconitate availability. **a**, Lipid synthesis pathways in *Y. lipolytica*. **b**, Intracellular total lipid concentration, lipid ratio as percentage of biomass, and maximum specific growth rates of the starting strain OKYL029 and lipid blocked strain JFYL007. **c**, Micrographs of *Y. lipolytica* cells stained with fluorescent dye Bodipy. **d**, Fatty acid methyl esters test for OKYL 029 and JFYL007. **e**, Cis-aconitate from medium availability in different strains. All data represent the mean of n = 3 biologically independent samples and error bars show standard deviation. ACC1, acetyl-CoA carboxylase; ACL, ATP-citrate lyase; ARE1, Acyl-CoA:sterol O-acyltransferase; DGA1, Acyl-CoA diacylglycerol acyltransferase 1; DGA2, Acyl-CoA diacylglycerol acyltransferase 2; FAS1 and FAS2, fatty acid synthase complex; LRO1, phospholipid: diacylglycerol acyltransferase; RAP, phosphatidic acid phosphatase; SCT1, glycerol-3-phosphate acyltransferase; SLC1, lysophosphatidic acid acyltransferase; TGL1, Cholesterol esterase; TGL3 Triacylglycerol lipase 3; TGL4, Triacylglycerol lipase 4.

**Fig. 3.**
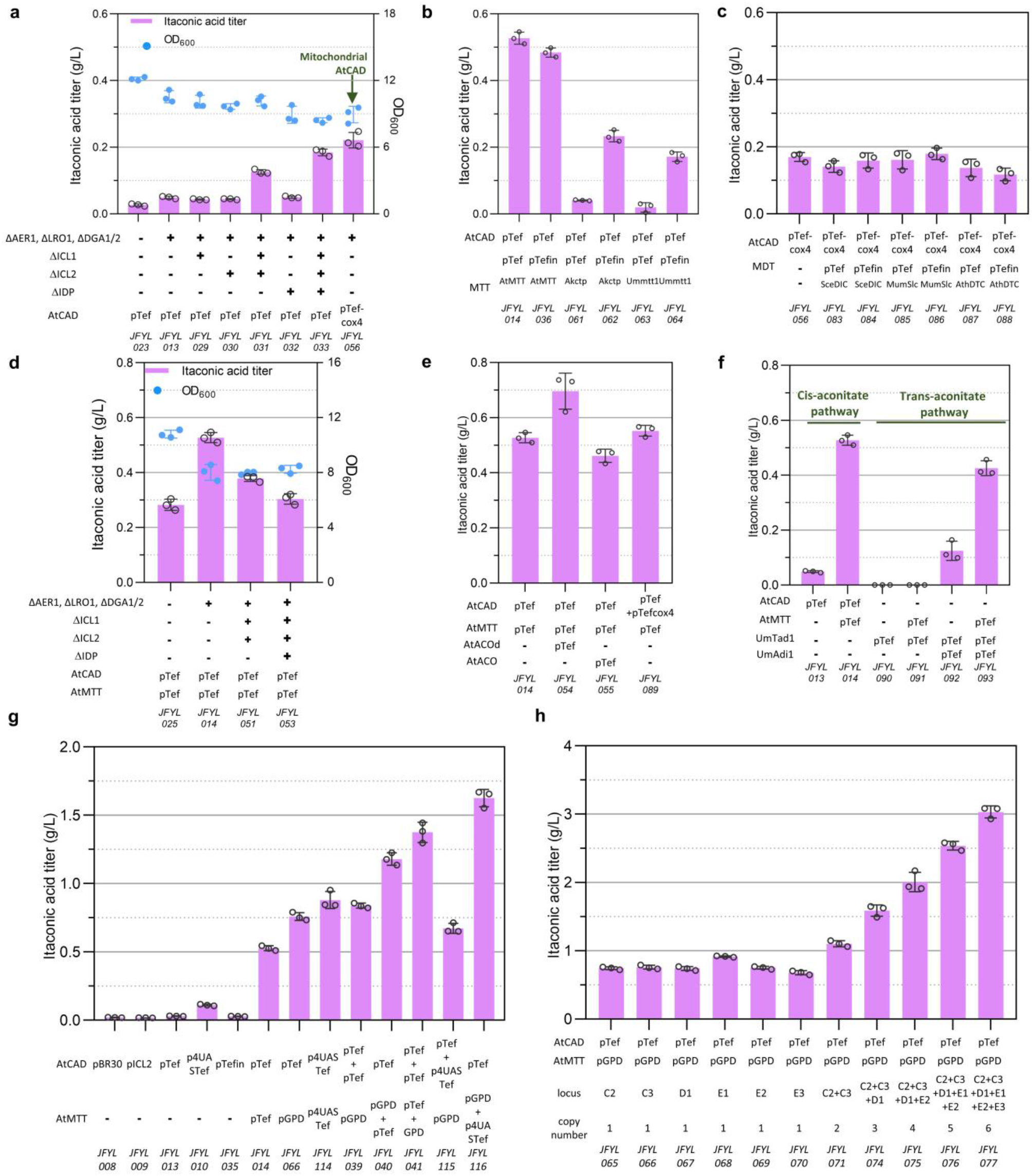
Optimize IA synthetic pathways. **a**, IA production with the introduction of AtCAD in IA sink pathways blocked strains. **b**, IA production in cytosol with 3 different MTT transporters under pTef and pTefin. **c**, IA production in the mitochondria with 3 different MDT transporters under pTef and pTefin. **d**, the combination of promising targets from module 1 and the selected AtMTT. **e**, introducing the ACOd expressed in cytosol increased the IA production, while combining the two IA synthetic pathways in cytosol and mitochondria resulted in unchanged IA production. **f**, the comparison of IA production through cis-aconitate and trans-aconitate. **g**, modulate expression of AtCAD and AtMTT under different promoters and their combination. **h**, increase the copy number of the selected combination resulted in increased IA production. All data represent the mean of n = 3 biologically independent samples and error bars show standard deviation.

Both the cytosolic and mitochondrial routes require transporters that we subsequentially studied. For cytosolic IA production, the tricarboxylic acid cis-aconitate is transported from the mitochondria. Contrastingly, mitochondrial IA production requires the produced dicarboxylic acid (IA) to be transported to the cytosol (Fig. 1). Therefore, three mitochondrial tricarboxylic acid transporters (MTT) and three mitochondrial dicarboxylic acid transporters (MDT) from different species were introduced under two strong promoters, pTef1 and pTefintron (17 times higher expression level than pTef1 ^45^). As shown in Fig. 3b, the AtMTT transporter from *A. terreus* under pTef1 promoter resulted in 0.527 g/L (JFYL014), which showed no significant difference with the mean IA titer by JFYL036 under pTefintron promoter, suggesting that extremely high expression of AtMTT is not beneficial for IA accumulation. Meanwhile, when expressing MTTs from *Aspergillus kawachii* (AkCTP) and *U. maydis* (UmMTT1), pTef1 gave almost no increase (Fig. 3b) over its parental JFYL013 strain (Fig 3a), while pTefintron yielded ca. five times higher IA production, suggesting low MTT activity of these two transporters. As for MDT transporters, none of the candidates, i.e. *Saccharomyces cerevisiae* (SceDIC), *Mus musculus* (MumDIC), and *Arabidopsis thaliana* (AthDIC), increased IA production (Fig. 3c). As a result, the cytosol pathway was selected for the next studies.

To further enhance AtMTT-facilitated cytosolic IA production, the abolition of sink pathways was tested. Blocking lipid accumulation yielded higher IA production when expressing CAD and MTT (JFYL014 in Fig. 3d), consistent with increased cis-aconitate production (JFYL007 in Fig. 2e). Unexpectedly, deletion of ICL1 and ICL2, which improved cis-aconitate and IA accumulation without the AtMTT transporter (Fig. 2e, Fig. 3a), resulted in decreased IA production (Fig. 3d), which was further exacerbated by IDP co-deletion. Therefore, only disruption of lipid accumulation (i.e. JFYL014) was retained in all later IA strains. Moreover, overexpression of mitochondrial AtACO that converts CA to cis-aconitate did not improve IA production (Fig. 3e), while cytosolic overexpression of AtACOd without mitochondrial leading sequence (MLS) increased the IA titer to 0.696 g/L, implying that considerable amounts of cytosolic CA are transformed to cis-aconitate for IA production. Attempts to facilitate tunnelling using a flexible GSG linker in a AtACO-AtCAD fusion protein did not result in improved production (Fig. S1). In addition, overexpression of both mitochondrial and cytosolic AtCAD (JFYL089, Fig. 3e) showed unchanged IA titers compared to sole cytosolic AtCAD.

Trans-aconitate derived IA production from *U. maydis* was then tested. No detectable IA accumulation was observed with expression of trans-aconitate decarboxylase (UmTad1, Fig. 3f), which is consistent with no detectable trans-aconitate level. Further introduction of UmAdi1 (JFYL092, Fig. 3f) afforded 0.125 g/L IA, which seemed very promising as the cis-aconitate pathway without the AtMTT transporter only resulted in 0.049 g/L IA (JFYL013 Fig. 3a). However, performance after introducing the AtMTT to the trans-aconitate pathway was not as good (0.426 g/L) as the cis-aconitate pathway (0.527g/L, Fig. 3f). Therefore, cytosolic AtCAD-based IA production with the AtMTT transporter remained the preferred strain.

The promising IA synthetic pathway was then fine-tuned by employing various promoters and copy numbers of AtMTT and AtCAD (Fig. 3g). Promoters pBR30, pICL1, pTef, p4UASTef and pTefintron with increased expression level were tested, and superior IA production was observed when using p4UASTef for AtCAD. After the introduction of AtMTT to afford a high cytosolic cis-aconitate availability, changing the AtMTT promoter from pTef to pGPD increased IA titers from 0.527 (JFYL014) to 0.759 g/L (JFYL066). Notably, the IA titer remained unchanged when introducing another copy of AtCAD under pTef (JFYL039), but decreased with an additional copy under the slightly stronger p4UASTef (JFYL115). On the contrary, the IA titer increased when another copy of AtMTT under pTef (JFYL040) or p4UASTef (JFYL116) was introduced. Interestingly, it seemed that higher AtMTT expression contributed more to IA production, while higher AtCAD levels alone seemed detrimental to production. As CAD was recently found to be toxic to the cell ^46^, these results indicate that the expression of AtMTT and AtCAD need to be finely tuned and matched. In consideration of the complex and repeated structure of the promoter p4UASTef, the combination of pTef for AtCAD and pGPD for AtMTT was instead selected for further study. When integrated at different genetic loci, this promising combination resulted in similar IA production (Fig. 3h), and significant increase of IA was achieved by introduction of multiple copy-numbers in JFYL077, where 6 copies of the pTef-AtCAD and pGPD-AtMTT combination afforded the best IA production. However, further overexpression of AtACOd did not show synergistic effect on IA production with 6 copies of the combination (Fig. S2), indicating a high flux through cis-aconitate.

### Mimic the regulation of nitrogen limitation

The IA producer JFYL066 (JFYL007 with one copy number of AtCAD and AtMTT) produced similar IA titer in both the NR and NL condition (Fig. 4a) when cultivated until glucose depleted (4 and 8 days, respectively). While titers were similar after glucose depletion, the NL condition resulted in higher IA yield and much lower productivity, probably due to the low biomass. It would therefore be promising to mimic the NL condition by partially imitate its regulation in NR conditions, to achieve both high yield and productivity.

**Fig. 4.**
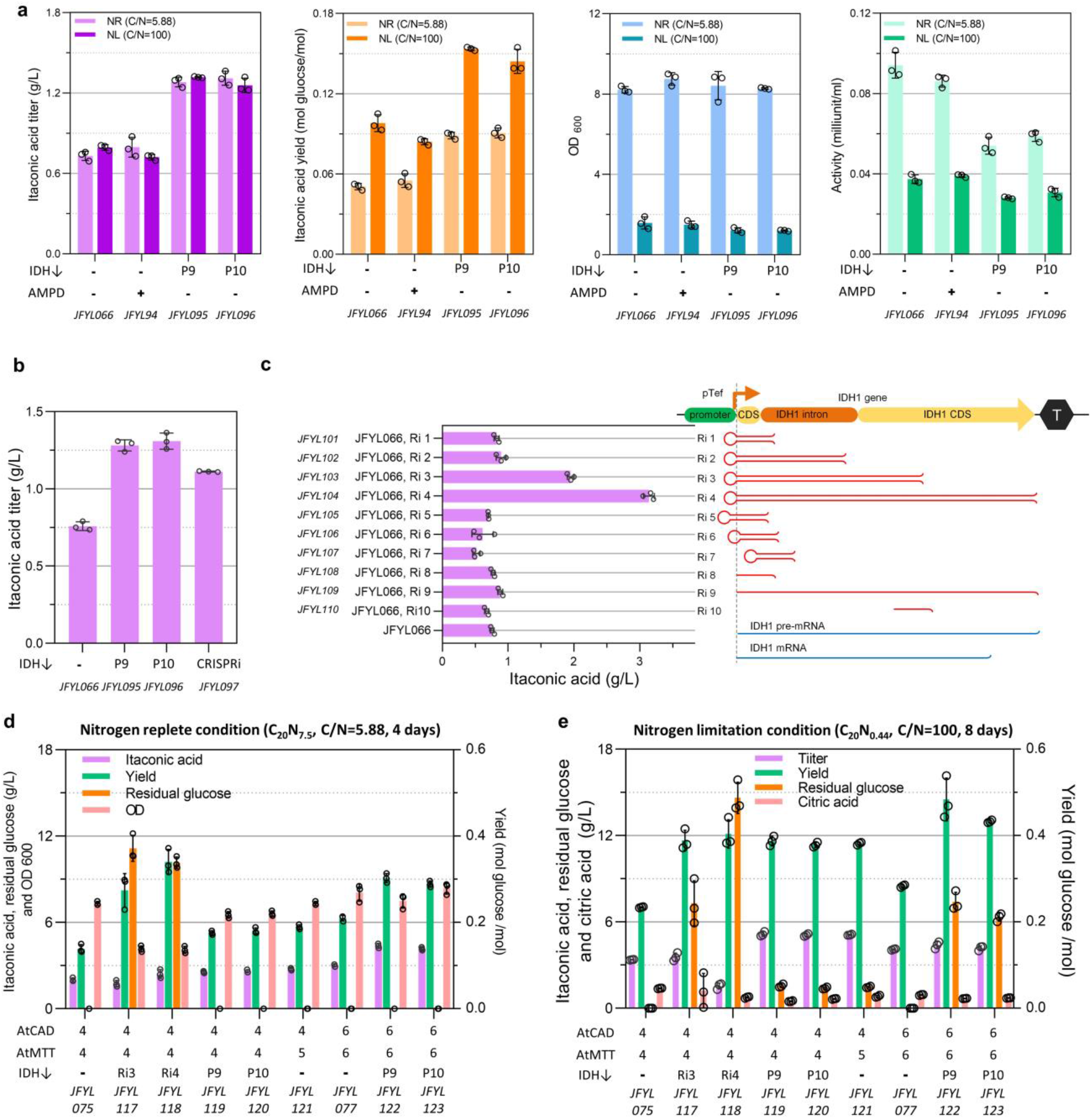
Mimic NL by down regulate IDH. **a**, the IDH was down regulated by overexpression of AMPD, and changing IDH’s native promoter P9 and P10. The IA titer, yield, OD_600_ and IDH activity were measured. **b**, comparison of the effect of IDH down regulation by promoter changing and CRISPRi. **c**, the effect of different RNA interference types on IA production. **d**, IA production in a NR condition. **e**, IA production in NL condition. All data represent the mean of n = 3 biologically independent samples and error bars show standard deviation.

It has been well reported that NL increases the activity of adenosine monophosphate (AMP) deaminase (AMPD) that converts AMP into inosine monophosphate (IMP), while IDH as AMP-dependent dehydrogenase is inhibited by this low AMP level ^47, 48^. Inconsistent with previous reports ^32^, overexpression of native AMP deaminase showed no positive effect on IA production (Fig. 4a). Next, IDH was down regulated by changing its native promoter for a weaker version, and 10 candidates were selected based on RNA seq data ^49^ (Fig. S3). As earlier attempts to delete the IDH ORF failed (Module 1), it also proved challenging to get positive transformants when the engineered promoter expression level was too low. Only two promoters, P9 and P10 were successfully exchanged. The IA titer and yield of these two strains were increased significantly in both NR and NL conditions, while the IDH activity and OD_600_ decreased (Fig. 4a). In general, IDH activities and IA yields anticorrelated in the NL conditions, but at the cost of prolonged incubation time and low productivity. Indeed, downregulation of IDH in NR conditions mimicked the positive effect of NL on IA production.

Because of the challenges encountered when downregulating essential *Y. lipolytica* genes using weak promoter exchange, we employed additional silencing methods to alternatively downregulate IDH, namely CRISPRi and RNAi. CRISPRi, allowing for sequence-specific repression of gene expression at the DNA level, was established by introducing a dCpf1 (Fncas12a D917A, E1006A) into *Y. lipolytica*^*50, 51*^. Four gRNAs were designed that would target the IDH subunit 1 coding gene simultaneously (Fig. S4), and the resulting strain produced 1.12 g/L IA (Fig. 4b), indicating that CRISPRi worked in *Y. lipolytica*. However, the IA titer was still lower than those with weak promoter exchanged (Fig. 4b). Alongside, we established RNAi^52^ in *Y. lipolytica* for the first time (Fig. 4c, S5) and prove it works. After exploring different silencing strengths by varying the length of shRNA or antisense ssRNA, strains with the longest shRNA (Ri3 and Ri4, Fig. 4d), which would be expected to have the strongest silencing, increased IA titer by 157% and 314% (1.95 g/L and 3.14 g/L), respectively.

The IDH downregulation strategy was then combined with modulation of AtCAD and AtMTT from Module 2. JFYL117 and JFYL118 afforded much higher IA yields than in JFYL075 (Fig. 4e). Unfortunately, excessive glucose was left in the RNAi strains, and therefore, JFYL118 in the 4-copy chassis showed lower IA titer (Fig. 4d). Meanwhile, JFYL119 and JFYL120 produced similar IA titer but lower yield due to its depletion of glucose compared with JFYL118. Moreover, increase the copy number of AtCAD and AtMTT to 6 copies could further increase the IA titer and yield, while JFYL122 resulted in the highest titer (4.3 g/L IA) with a yield of 0.31 mol IA/mol glucose. When NL condition (Fig. 4f) was employed, all strains exhibited much higher IA yields but lower productivities. Notably, IA with similar titer and yield was produced by JFYL122 in NR condition (4 days) and JFYL077 in NL condition (8 days), indicating that we successfully mimicked the NL with enhanced productivity by downregulation of IDH and increased the productivity by 2-fold.

As such, JFYL122 with 6 copies of AtCAD and AtMTT, and P9 promotor modulated IDH expression was selected for further study, due to its superior titer in NR condition and highest yield in NR condition, compared with all IDH down-regulated strains.

To present the different flux in NL and NR conditions, flux balance analysis (FBA) was performed by constraining with measured exchange reaction rates and modified biomass compositions from chemostat bioreactor cultivations (Fig. 5). The predicted flux in JFYL007 indicated that the flux of TCA is higher from NL condition than that from NR condition. The higher flux could increase cis-aconitate availability, which contributed to high IA yield. Therefore, our strategy to decouple NL and high flux in TCA by mimic NL regulation, allowing higher biomass, could afford balanced IA production with both high yield and productivity. Notably, this simulation especially contributes to IA production when NL should be avoided in Module 4 in the next section.

**Fig. 5.**
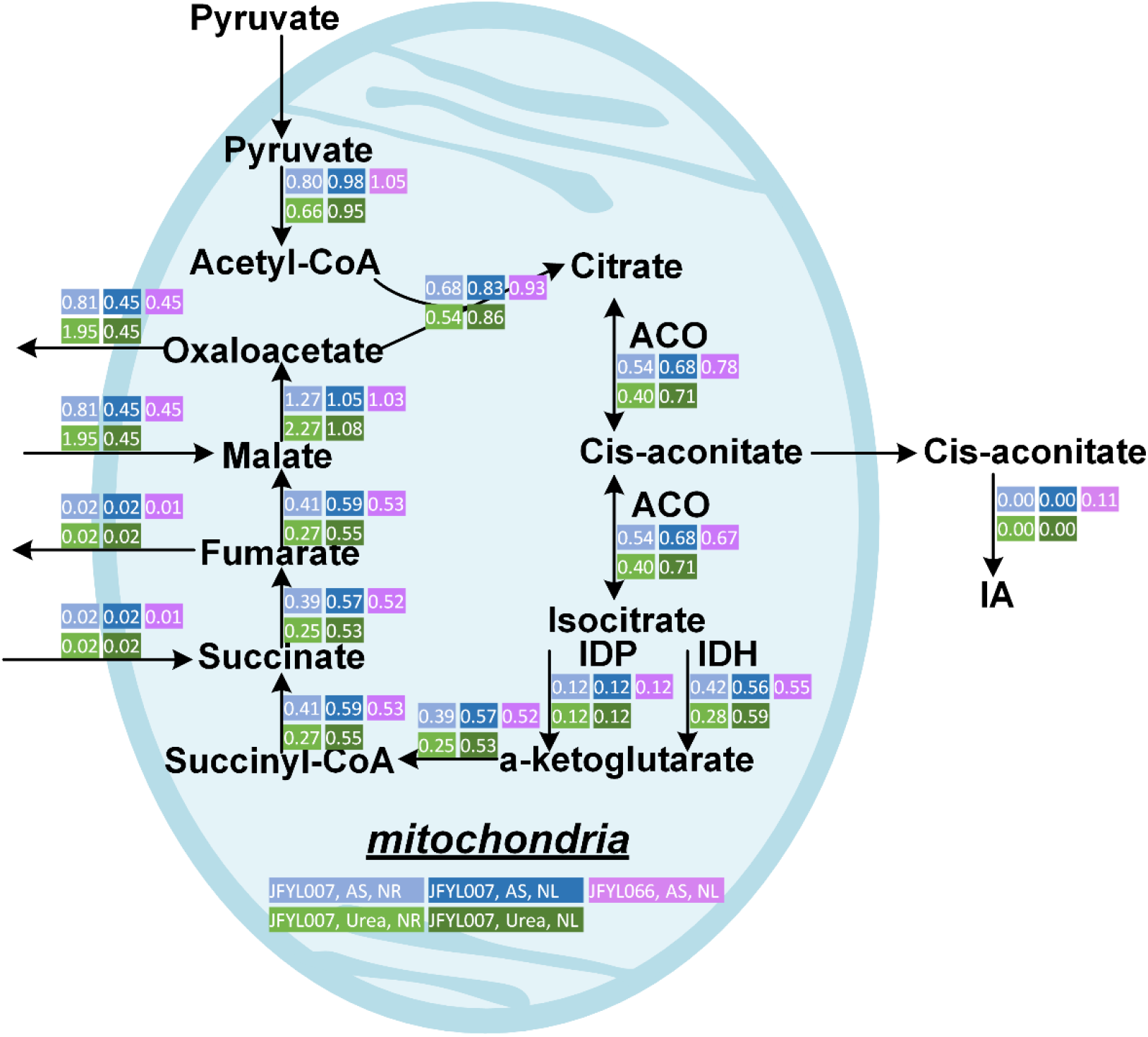
Flux balance analysis of JFYL007 and JFYL066 in nitrogen replete and nitrogen limitation conditions. Strains were cultivated in modified Delft medium with different nitrogen source (ammonia sulfate or urea) and absolute nitrogen amount. NR nitrogen replete, NL nitrogen limitation.

### Fermentation engineering and scale-up

The most promising strain (JFYL122) was cultivated in 1L fed-batch bioreactors for scaling up from flask level. As NL could support higher yield, we first tested a widely used nitrogen control strategy which provided NR condition for high biomass initially and then generated NL conditions by the consumption of the nitrogen source for IA production with enhanced productivity. Unexpectedly, significant amounts of CA started to accumulate after three days (Fig. 6a), and soon surpassed the IA titer, with a final titer of 67.5 g/L and 17.1 g/L, respectively. The predominant CA accumulation (Fig. 6a) was possibly due to the high complex of NL regulation where both absolute nitrogen amount and C/N ratio would affect metabolism. While glucose was fed during the cultivation, nitrogen likely became limiting at day 3, when growth halted at OD_600_ 120. At this point, abundant CA started to be secreted, indicating that it was not being converted to cis-aconitate. Contrastingly to what we measured before, only ca. 1 g/L CA was produced in NL condition in deep-well plates (Fig. 4f). As scale up conditions were not directly transferable, further fermentation engineering is required to optimize production in bioreactors.

**Fig. 6.**
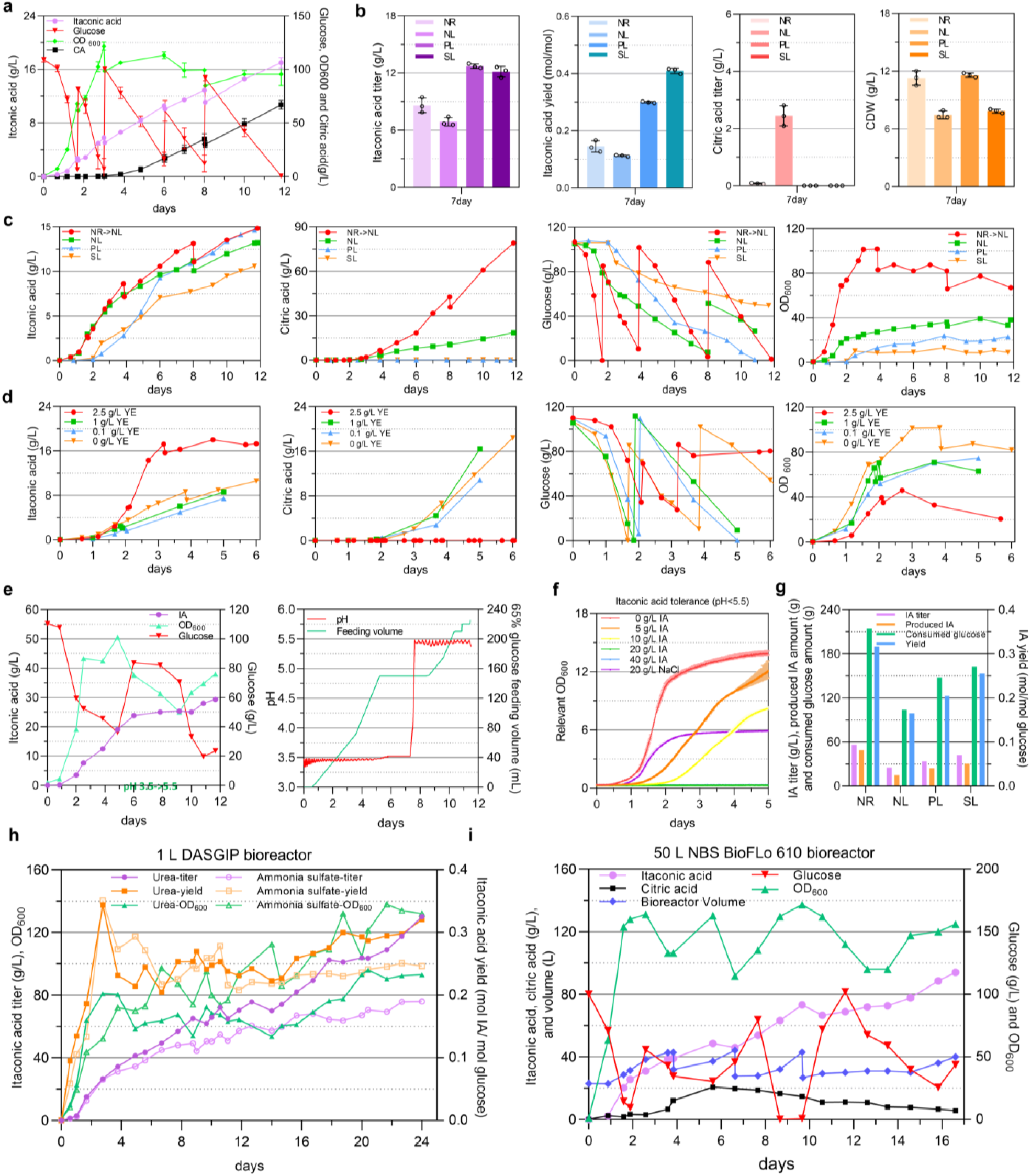
IA production in bioreactors. **a**, JFYL121 was cultivated in 1L bioreactors in Delft medium with 10 g/L ammonia sulphate (initial C/N=22, 65% glucose was fed in pulse form). Data was from a representative run. **b**, JFYL121 was cultivated in flasks for batch under nitrogen replete (NR, C_100_N_10_), nitrogen limitation (NL, C_100_N_2.5_), phosphate limitation (PL), and sulfur limitation (SL) conditions. All data represent the mean of n = 3 independent samples. **c**, JFYL121 was cultivated in 1L bioreactors under nitrogen switch (initial C_100_N_10_, from NR to NL), nitrogen limitation (NL, C_100_N_2.5_), phosphate limitation (NL), and sulfur limitation (SL) conditions. **d**, the effect of addition of yeast extract on IA titer, CA accumulation, glucose consumption and OD_600_ in C_100_N_10_ (C/N=22) delft medium. **e**, itaconic acid tolerance in Delft medium in biolector. **e**, JFYL121 was cultivated in 1L bioreactors, and continuous feeding of glucose and YE resulted in enhanced IA production. The pH was changed from 3.5 to 5.5 after the cell growth stopped, and then the strain began to produce IA again. **f**, IA tolerance measured in biolector. **g**, JFYL121 was cultivated in 1L bioreactors under nitrogen replete, nitrogen limitation, phosphate limitation, and sulfur limitation conditions with the continuous feeding of glucose and YE together and pH was set at 5.5. **h**, JFYL121 was cultivated in 1L bioreactor at bench-top scale, and AS and urea were used as the nitrogen source, respectively. **i**, JFYL121 was cultivated in 50L bioreactor at semi-pilot scale.

To reduce CA accumulation, phosphate limitation (PL) and sulphur limitation (SL) were evaluated. JFYL122 was tested on different cultivation media in flasks with higher glucose (100 g/L) than tested previously (20 g/L, Fig 4e-f). CA only accumulated if nitrogen was the limiting nutrient (NL-bioreactor medium), while both PL and SL gave higher IA titers and yields (Fig. 6b). These results were further tested in bioreactors (Fig. 6c), and only cultivation in PL and SL precluded CA production, yielding higher IA yields at the cost of low glucose consumption rate (Fig. S6). These above results further support nitrogen limitation should be avoided to eliminate CA secretion, and this should likely be accomplished by adding nitrogen source to support higher OD_600_ and not alternative limiting nutrients that yields lower productivity.

To increase IA production, these following modifications were tested. For higher productivity, as yeast extract (YE) with low production cost is frequently used to improve cell growth^53^, different concentration of YE were tested in NL-bioreactor medium. 2.5 g/L YE significantly increased the IA titer to 17.3 g/L (Fig. 6d) within 3 days. With lower YE supplementation, CA again started to accumulate from the second day, indicating depletion of the nitrogen source. Meanwhile, as high osmotic pressure is reported as a key factor responsible for inhibition of yeast growth and decrease fermentation performance^54^, changing from pulsed feeding to continuous feeding could reduce potential osmotic stress effects, which indeed significantly increased IA production to 24.5 g/L at the sixth day (Fig. 6e). However, after that, the cells suddenly arrested glucose consumption and IA production. Next, changing the pH from 3.5 to 5.5 initiated the cells to recover and continue to produce IA (Fig. 6e). Indeed, testing IA tolerance in high-through put microbioreactors with an initial pH of 3.5 showed that 20 g/L IA significantly interrupts cell growth (Fig. 6f), while it was reported that *Y. lipolytica* is highly tolerant to 60 g/L IA at pH 7 ^33^. Therefore, addition of YE, continuous feeding, and pH of 5.5 were the key factors in later cultivations.

The above optimized conditions were then combined, and no growth disruption was observed when IA titer was above 25 g/L with pH controlled at 5.5 (Fig. 6g). 600 g/L glucose with 20 g/L YE were continuously fed together. As anticipated, the PL and SL conditions afforded better IA production than the NL condition (Fig. 6g). Significantly, NR condition afforded higher IA productivity, as well as titer and yield, indicating the growth associated strategy with unlimited nitrogen source and the mimic of down regulation of IDH works well here. This condition was further test in bioreactors and 68.1 g/L IA with 16.75 days (Fig. 6h) with ammonia sulfate as the nitrogen source. However, to maintain pH at 5.5, base (6 M KOH) supplementation with large volume was required (Fig. S7). This drastically increased the cultivation volume during this period, resulting a fluctuation of the IA titer after 17 day, although the absolute amount of IA still increases (Fig. S6). Notably, the consumption of ammonium sulfate reduces pH while the use of the alternative and cheap nitrogen source urea would not acidify the media^55^. Alternatively using urea reduced the base feeding volumes and significantly increased IA titer to 130.5 g/L IA with a yield of 0.320 mol/ mol glucose. These results suggested the successfully establishment of our engineered strain as promising candidate for IA production.

To verify the performance of JFYL122 during the scaling up at semi-pilot scale, a 50L bioreactor was employed with urea as the nitrogen source. Within 1.5 days, the OD_600_ quickly rose to 160 and the highest productivity was obtained before day 9.17. Due to accidental glucose limitation at that time, the cells lost their vitality and it took them 3 days to recover. However, the IA production then continued with high productivity. A final 94.1 g/L IA with and a productivity of 0.238 g/L/h and a yield of 0.36 mol/ mol glucose was reached after day 16.58. The total amount of IA produced during this fermentation was 6161.9 g. These results indicated that our engineered strain exhibited robustness and potential during the scale up process, indicating it is promising IA producer at industrial scale fermentation.

## Discussion

Through a modular approach to rewiring *Y. lipolytica*, we gradually enabled and increased IA production from 0.027 to 130.5 g/L. Notably, in the strain development process, some strategies only have positive effect on IA production at certain conditions. When trying to further increase the production, these strategies may not be beneficial or not synergistic when combined. This could be explained by the complexity of the metabolism in *Y. lipolytica*, demonstrating the difficulty for enhancing production and scale-up. Since our goal was to obtain enhanced IA production in *Y. lipolytica*, and transform it to industrial level production, the final combination of them were carefully tested to balance the titer, yield and productivity.

We first tried to increase cis-aconitate availability by decreasing flux of sink pathway from CA and ICA nodes in module 1. CA derived lipid accumulation was significantly reduced, and the increased cis-aconitate level and IA production (Fig. 3a) suggest successful redirection of its flux to IA production. This is thereby the first report to demonstrate that blocking all four acyltransferases in lipid accumulation can afford enhanced production of desired product in *Y. lipolytica*. For the ICA node, deletion of both ICL1 in glyoxylate cycle and ICL2 in methylcitrate cycle increased the cis-aconitate availability in the cytosol, which had a positive effect on the IA titer after CAD introduction (Fig. 3a). However, with the increased availability of cis-aconitate after the introduction of AtMTT transporter, the deletion of ICL1/2 had a negative effect on the IA titer. Note that the isocitrate lyase (ICL1) reaction was almost inhibited by IA higher than 0.13 g/L ^56^, while both ICL1 and ICL2, act on 2-methylisocitrate ^57, 58^, are not inhibited by IA. Therefore, we speculate that interrupt of 2-methylisocitrate cycle could have a negative effect on IA production with higher IA concentration above 0.26 g/L. Notably, there is conflicting information about the function and location (mitochondria or cytosol) of ICL2, and the effect of its deletion on IA production need to be further studied. Meanwhile, two isocitrate dehydrogenases, IDP and IDH, are interrupted for the ICA node. Deletion of the NADP^+^ dependent IDP slightly increased the cis-aconitate level, but the IA titer was almost unchanged (Fig. 3a). This might be explained by its low activity, while in *Y. lipolytica*, NADPH for lipid synthesis is mostly provided by the phosphopentose pathway, other than the pathway involved by IDP^59^. The deletion of the essential gene IDH was not achieved, while its downregulation was carefully studied in Module 3.

In module 2, introduction of mitochondrial tricarboxlic transporters increased IA production significantly, due to increased cis-aconitate availability. A new IA synthetic pathway form trans-aconitate was tested and was likely dominant at low cis-aconitate and IA levels, where twice amount of IA was produced by JFYL092 with TAD than JFYL013 with CAD. However, after introducing AtMTT, JFYL093 with trans-pathway produced less IA than JFYL014 with cis-pathway, possibly due to the two steps convertion and a possible low kinetics parameters of TAD, while AtCAD exhibited a high Kcat and Km ^60^. This phenomenon also observed elsewhere that trans-pathway led to an IA titer increase at low IA level due to the high proportion (88%) of trans-isomer of aconitate at equilibrium, while produced slightly lower IA amount at a high IA level due to unutilized substrate ^46^. Indeed, fine modulation of cis-pathway and increased its flux significantly increased IA production.

In module 3, without nitrogen limitation, the NL regulation was mimicked by IDH knockdown to afford relatively high IA yield from NL, and retain high productivity from NR. To enable downregulation of essential genes such as IDH, we tested promoter replacement, and established RNAi and CRISPRi. We believe it is the first time to employ RNAi in *Y. lipolytica*, and the first time to co-employ CRISPR/cas9 for gene editing and CRISPRi/dcas12a, which could significantly increase the genetic toolbox in *Y. lipolytica* for gene knockdown. Therefore, the gene knockdown technology developed here expands the gene toolbox of *Y. lipolytica*. Significantly, the regulation of NL was successfully mimicked in NR condition (Fig. 4e, 4f). However, to afford a synergetic effect with enhanced IA synthetic pathway in Module 2, a changed weaker promoter P9 preformed best to balance cell growth and IA productivity. Furthermore, the down regulation of IDH could even further increase the IA yield in NL condition (Fig. 4a, 4e, 4f), suggesting a deep repression than previously widely used NL, which could be employed to enhance CA or lipids production in *Y. lipolytica*.

During the scale-up process in module 4, excess CA secretion with NL, and IA tolerance became main challenges which was not exhibit at flask level. To decrease CA secretion, four limitation conditions (NL switch, NL, PL, and SL) were compared (Fig. 5b, 5c) at both flask level and scale-up level. Notably, in *A. terreus*, PL was employed for IA production, while IA titer was decreased with NL ^22, 61^. Therefore, NL was be avoided in this study for *Y. lipolytica* to prevent CA secretion. Interestingly, CA was reported to be accumulated in all NL, PL and SL conditions, while here its secretion was totally abolished in PL and SL, which afforded much higher IA yield. However, due to low productivity, all nutrition limitations were avoided in this study. Meanwhile, for IA tolerance, it was reported that at pH 7 in YPD medium, *Y. lipolytica* is highly tolerant to itaconic acid and remains growth with 60 g/L IA^33^. However, here in Delft medium with pH below 5.5, cell growth was inhibited and could be recovered by adjusting the pH at 5.5 (Fig. 5e, 5f). Interestingly, when the high CA producer *A. niger* was engineered for IA production, its growth was hampered with 10 g/L IA, while the IA toxicity could be alleviated by maintaining nitrogen availability ^31^. This could be explained by the glutamate-dependent acid resistance mechanism (GDARM)^30^, which was employed by acid-tolerant microorganisms to reduce acids toxicity in transformed to γ-aminobutyric acid (GABA) to ultimately increase the alkalinity of the cytoplasm^30^. Furthermore, to maintain pH at 5.5, the volume increased significantly, and the medium had to be tapped out to not excess the maximum working volume, which however, could be avoided by starting with a lower volume and feeding ammonia instead. In our previous report, urea is a drop-in nitrogen source alternative to ammonia sulfate in *Y. lipolytica*, and there’s no significant concerted changes in the transcriptome^55^. Notably, without down regulation of IDH, the FBA based on measured exchange reaction rates from different conditions from chemostat (Fig. 6) shows that in NR condition, the flux through cis-aconitate by using urea is lower than that using ammonia sulfate, while in NL condition the cis-aconitate flux is similar when using both urea and ammonia sulfate. However, by mimic the regulation of NL, the IA titer, yield, and productivity are all higher when using urea compared to ammonium sulfate (Fig. 6h). Ultimately this proof-of-concept semi-pilot run suggested our engineered strain could producing IA with high titer. With improved fermentation control, the IA production could be further improved at this scale, suggesting our engineered *Y. lipolytica* as a competitive alternative to *A. niger*.

## Methods

### Strains, media and cultivation

*E. coli* strain DH5α was used for plasmid construction, which was grown at 37 °C with shaking in Luria-Bertani (LB) Broth (Sigma) supplemented with 50 μg/ml of ampicillin for plasmid propagation. All *Y. lipolytica* strains constructed in this study were derived from ST6512, which is a W29 background strain (Y-63746 from the ARS Culture Collection, Peoria, USA; a.k.a. ATCC20460/ CBS7504) and harbors *Cas9* at *KU70* locus for marker-free genomic engineering by EasyClone YALI toolbox^45^. The complete strains used in this study are listed in Supplementary Table 1.

Y. lipolytica strains were cultivated in the minimal mineral, a modified Delft medium^62^ with 0.2 M phosphate buffer, in shake flasks and 24 deep well plates unless stated otherwise. The medium contained 7.5 g/L (NH_4_)_2_SO_4_, 19.2 g/L KH_2_PO_4_, 10.2 g/L K_2_HPO_4_, 0.5 g/L MgSO_4_•7H_2_O, 2 mL trace metals solution stock, and 1 mL of vitamin solution stock. The trace metals and vitamin solution were added after autoclave. Trace metal solution contained 3.0 g/L FeSO_4_•7H_2_O, 4.5 g/L ZnSO_4_•7H_2_O, 4.5 g/L CaCl_2_•2H_2_O, 1 g/L MnCl_2_•4H_2_O, 300 mg/L CoCl_2_•6H_2_O, 300 mg/L CuSO_4_•5H_2_O, 400 mg/L Na_2_MoO_4_•2H_2_O, 1 g/L H_3_BO_3_, 100 mg/L KI, 19 g/L Na_2_EDTA•2H20. The vitamin solution stock contained 50 mg/L d-biotin, 1.0 g/L D-pantothenic acid hemicalcium salt, 1.0 g/L thiamin-HCl, 1.0 g/L pyridoxin-HCl, 1.0 g/L nicotinic acid, 0.2 g/L 4-aminobenzoic acid, 25g/L myo-Inositol. The initial pH was set at 6.5 with KOH. C_x_N_y_ was used here to indicate the glucose and nitrogen (ammonia sulfate or urea) amount (g/L) with x and y values. C/N was used to indicate the Carbon and Nitrogen molar ratio. In module 1 to 3, 20 g/L glucose and 7.5 g/L ammonia sulfate were used in this medium to set the NR condition (C/N=5.88), while in module 3, NL condition (C/N=100) was achieved with 20 g/L glucose and 0.44 g/L ammonia sulfate. In module 4, the media with varying C/N ratios, PL, and SL conditions were provided in Supplementary Table 5 to study the carbon flux distribution between CA and IA for optimal IA fermentation. For all bioreactor fermentation,

In chemostat cultivations, all modified Delft media contained 3 g/L KH_2_PO_4_, 0.5 g/L MgSO_4_•7H_2_O, 1 mL trace metals solution stock, and 1 mL of vitamin solution stock. For NL condition (C/N ratio 116), the media contained 0.471 g/L of ammonium sulphate or 0.213 g/L urea (sterile filtered) and 27.5 g/L glucose, while for NR condition (C/N ratio 3) the media contained 5.28 g/L of ammonium sulphate or 2.4 g/L urea (sterile filtered), 7.92 g/L glucose. pH was set at 5.5 with 2M KOH, and the temperature was set at 28°C.

All *Y. lipolytica* strains were cultivated at 30 °C for shake flask and 24 deep well plates fermentation with a shaking speed of 220 rpm. The 24 deep well plates are purchased from Enzyscreen (plates, CR1424a; lips, CR1224f), and the volume of culture is 2.5 mL.

### Cloning and transformation procedures

The complete list of plasmids and primers used in this work is available in the supplementary file S2 and S3. Plasmids for genome engineering were constructed with set of vectors from EasyCloneYALI as backbones [49]. Construction of plasmid for genome engineering was performed according to EasyCloneYALI instructions.

For gene deletion of ARE1, DGA1, DGA2, LRO1, ICL1, ICL2, IDH1, IDH2, and IDP, equal amounts of two single-stranded oligonucleotides (90-120 bp,100 pmol/μL) were used to obtain the repair templates. The pair of oligonucleotides were incubated for 5 min at 98°C and allowed to cool down to room temperature. The EasycloneYALI toolbox was used to construct gRNA plasmids for CRISPR/Cas9 aided gene deletion.

For gene overexpression, the integration plasmids were constructed with different promoters and genes (Supplementary Table 2). The marker-free integration cassettes were obtained by digesting the corresponding plasmids (Supplementary Table 2). After purification, around 1 µg of repair fragment was used for transformation. For each transformation around 500 ng of gRNA plasmid was used.

For IDH1 promoter change, the repair fragments were obtained by overlap PCR of the upstream of IDH1, promoter candidate, and downstream of IDH1. AKEC8 was used as the gRNA helper plasmid for IDH1 promoter change.

### Chemostat experiments

Chemostat cultivations were performed at 28°C in DasGip systems with 1 L stirrer -pro vessels (Eppendorf) as previously reported^63^. Briefly, the working volume was of 500 mL kept constant using an overflow pump. and pH was maintained at 5.0 ± 0.1 by automatic addition of 2M KOH. The agitation was set at 600 rpm, and aeration was maintained with sterile air at 30Lh-1 (1 vvm) and monitored with DO probes (Mettler). Delft medium was used in batch, and constant feed with a dilution rate of 0.10 h^−1^ was initiated after the exponential growth phase to obtain steady-state cultivation. The growth was monitored by the CO_2_ exhaust gas. Samples for lipid analysis from JFYL007 and OKYL029 were taken after 5 residence times of steady-state growth. Each condition was cultivated at least in triplicates.

### Lipid body visualization

Lipid body visualization was performed by microscopic analysis. Bodipy® Lipid Probe (2.5 mg/mL in ethanol, Invitrogen) was added to the cell suspension (OD_600_ of around 1) with 10 min incubation at room temperature. Zeiss Axio Imager M2 microscope (Zeiss, Le Pecq, France) with a 100× objective and Zeiss filters 45 and 46 for fluorescence microscopy were used to acquire the images.

### Lipid Extraction and Quantification

Samples of OKYL029 and JFYL007 from chemostat were harvested for measuring intracellular lipid concentration, lipid ratio. Fatty acids were extracted, transformed to fatty acid methyl esters (FAMEs), and quantified by GC-MS using a published method. GC-MS (Thermo Scientific Trace 1310 coupled to a Thermo Scientific ISQ LT) and a ZBFAME column (Phenomenex, 20 m, 0.18 mm Inner Diameter, 0.15 μm Film Thickness) were used for the quantification. The Fatty acids (C16:0, C16:1, C18:0, C18:1, and C18:2) were considered to calculate intracellular FAME amount, intracellular lipid concentration (total FAME amount per cell dry weight, mg fatty acid/mg dry biomass).

### High-performance liquid chromatography (HPLC) Analysis

Glucose and other extracellular metabolites in medium were quantified by HPLC UltiMate^®^ 3000 HPLC system (Dionex). Fermentation samples were harvested (samples with high OD from bioreactors were diluted by 10 times), centrifuged for 2 min at 13000× g, and the supernatant was used for HPLC analysis. Two HPLC systems was used. Luna 5µ C18(2), 250*4.6mm (Phenomenex) was utilized for testing cis-+aconitate. Its mobile phase was composed of 0.02 mM Monopotassium phosphate buffer (pH 2.65) to methanol (a 95:5 ratio of) with a flow rate of 1 mL/min. Aminex HPX-87H ion exclusion column (Bio-Rad, Solna, Sweden) was utilized for testing glucose, IA, CA and other metabolites. 5 mM H_2_SO_4_ was used as mobile phase with a flow rate of 0.6 mL/min. Glucose was quantified using a refractive index detector (Shodex, Munich, Germany) and other metabolites were quantified using a UV detector.

### FBA analysis

The *Yarrowia lipolytica* GEMs used in this work was refined based on a published GEMs named iYali v4.1.1^64^, during which the flux bounds for reactions y102884, y102948, y000303, y002305, y000659 and y000661 were constrained at zero to reflect the gene deletion and the manual protein compartment annotation. In addition, four new reactions were added into GEMs to represent the whole molecular process in synthesis and secretion of itaconate. During the simulation through a python package cobrapy, the mean flux values of the physiological parameters, including the growth rate, production rate and by-product formation rate were used to constrain GEMs. Afterwards, the minimization of glucose uptake rates was set as the optimizing objective function to solve the fluxes through each reaction based on the Parsimonious FBA (pFBA) procedure^65^.

### Bioreactor fermentations

Bioreactor fermentations at bench-top level were performed in DasGip 1-L stirrer-pro vessels (Eppendorf, Jülich, Germany) at 30 °C. pH was monitored with a pH sensor (Mettler Toledo, Switzerland) and maintained at set value by automatic addition of 6M KOH or 2M HCl. Bioreactor fermentations at semi-pilot level were performed in New Brunswick BioFlo 610 bioreactor systems.

### IDH assay

IDH enzyme activity was determined by the IDH activity detection kit (#MAK062, Sigma, USA), according to manufacture’s instruction of the assay kit.

### IA tolerance experiment

Modified Delft medium (with addition of MES to a final concentration of 0.2M to reduce the influence of pH change) was supplemented with various concentrations of IA (0, 5, 10, 20, 40 g/L) or 20 g/L NaCl as a negative control to test the IA tolerance. The initial pH was 6.5. The strains were cultured in 48 deep well plates in microbioreactors Biolector at 30 °C with the initial OD_600_ of 0.2. The OD_600_ was measured every 30 minutes automatically.

## Author Contributions

Conceptualization: E.J.K. and J.F.; investigation, J.F., S.Z., H.L., O.K., N.P., A.K., and D.K.; writing— original draft, J.F.; writing—review and editing, J.F., S.Z., O.K., N.P., and E.J.K.; supervision, E.J.K. All authors have read and agreed to the published version of the manuscript.

## Acknowledgement

This research was funded by the Novo Nordisk Foundation [grant NNF10CC1016517], the Research Council for Environment, Agricultural Sciences, and Spatial Planning (Formas) [grant 2018-00597], and the Swedish Research Council (VR) [grant 2019-04624]. We thank the BRISK2 [European Union’s Horizon 2020 Research and Innovation Programme under grant agreement 731101] and COST Action YEAST4BIO (CA18229) to support the semi-pilot scale fermetation experiment.

## Notes

### Competing Interest Statement

The authors have declared no competing interest.

