## Supplementary figures and tables for "Reprogramming *Yarrowia lipolytica* metabolism for efficient synthesis of itaconic acid from flask to semi-pilot scale"

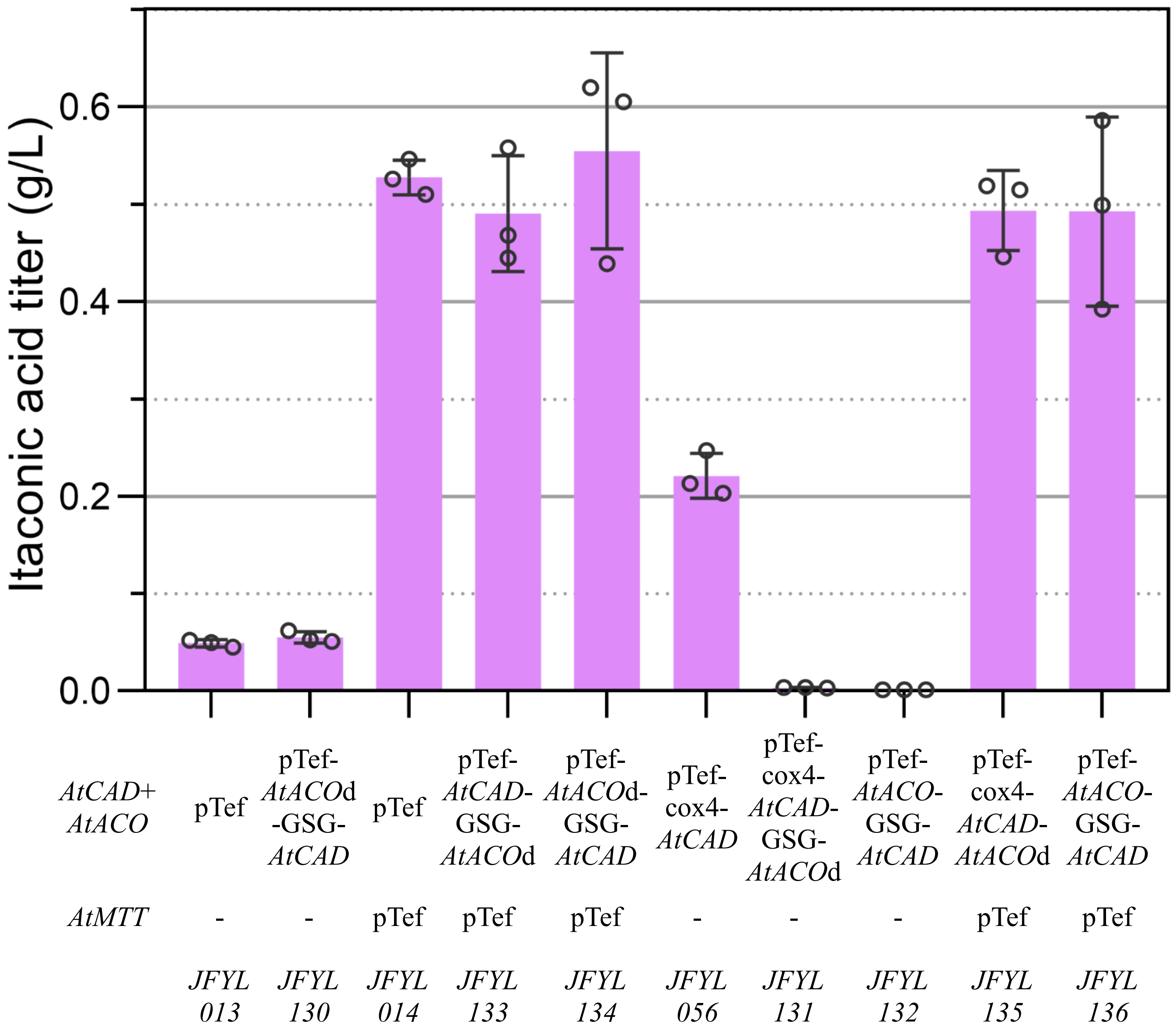


Fig. S1 *AtACO* and *AtCAD* fusion proteins with flexible linker GSG

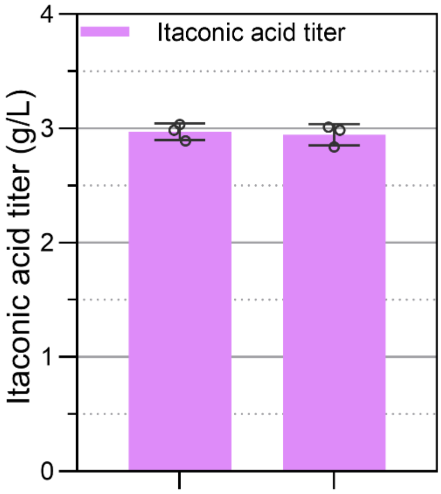


| | | |
|---|---|---|
| *AtCAD* | pTef | pTef |
| *AtMTT* | p*GPD* | p*GPD* |
| Copy number | 6 | 6 |
| *AtACO*d | - | pTef |
| | JFYL128 | JFYL129 |

Fig. S2 IA production in JFYL128 and JFYL129

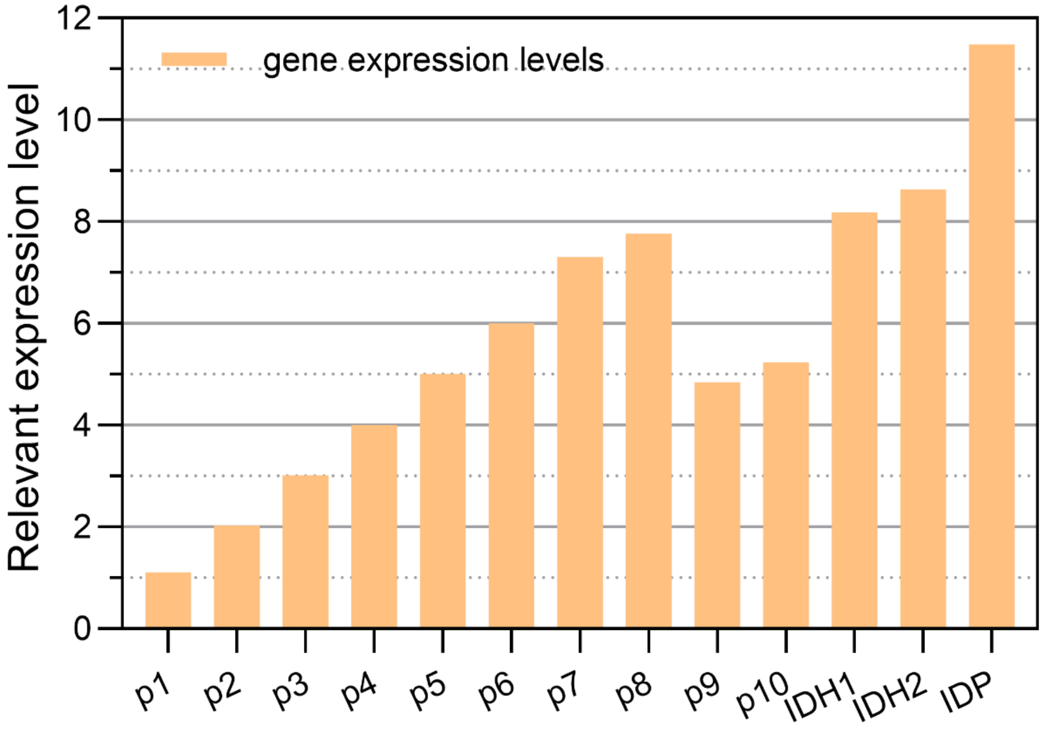


| Gene | Gene | Mean WTN | Note |
|---|---|---|---|
| p1 | YALI0A00550g | 1.1 | Stable expression |
| p2 | YALI0D10967g | 2.03 | |
| p3 | YALI0B16192g | 3.01 | |
| p4 | YALI0F20680g | 4.01 | |
| p5 | YALI0E13596g | 5.01 | |
| p6 | YALI0B07667g | 6 | |
| p7 | YALI0D05731g | 7.30 | Decreased expression at high CN ratio |
| p8 | YALI0C16885g | 7.76 | |
| p9 | YALI0F03344g | 4.84 | |
| p10 | YALI0A14388g | 5.23 | |
| *IDH1* | YALI0D06303g | 8.18 | |
| *IDH2* | YALI0E05137g | 8.63 | |
| *IDP* | YALI0F04095g | 11.48 | |

Fig. S3 Gene expression level of candidates for weaker promoter changing.

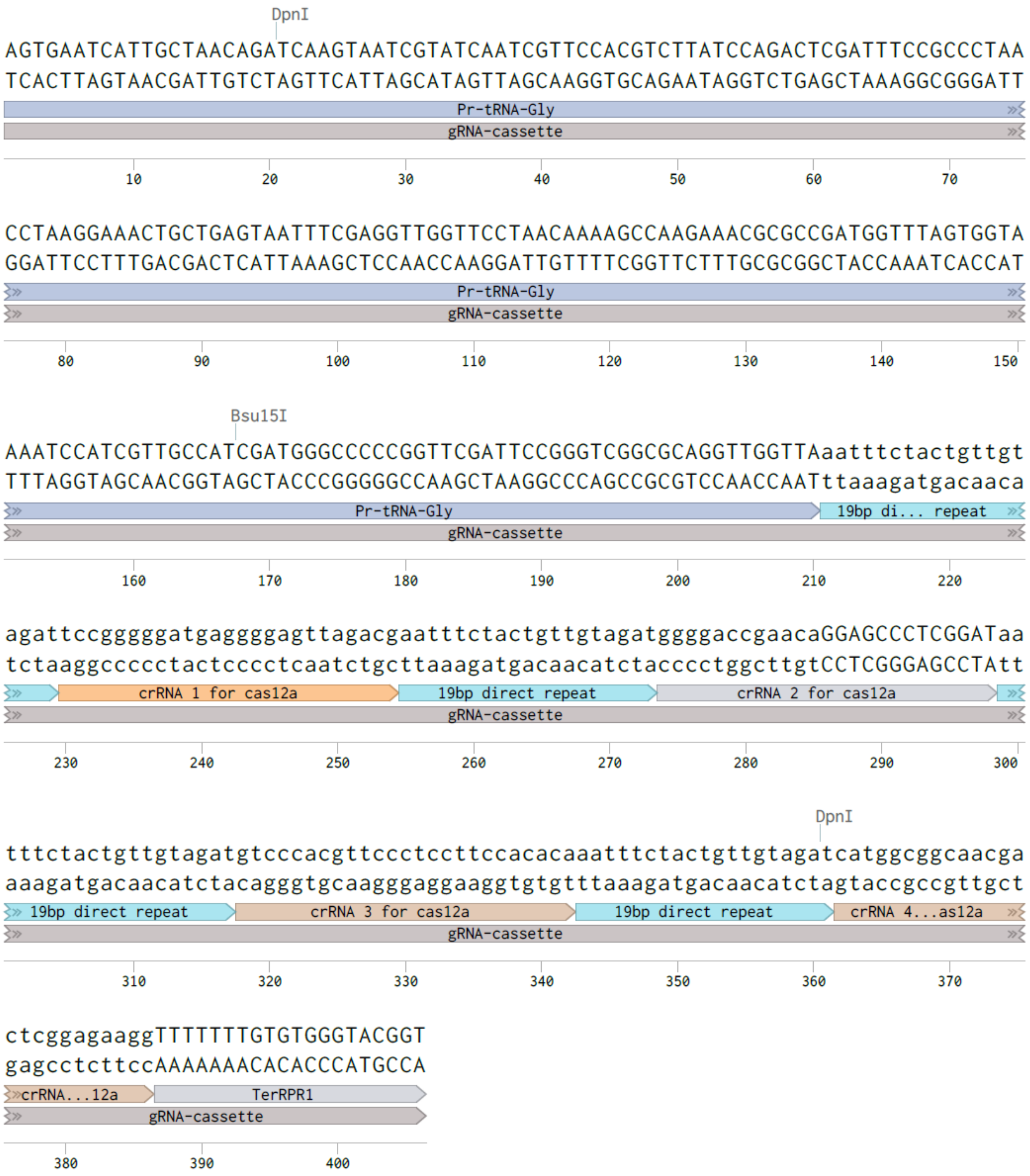


Fig. S4 4 gRNA targeting to *IDH1* CDS

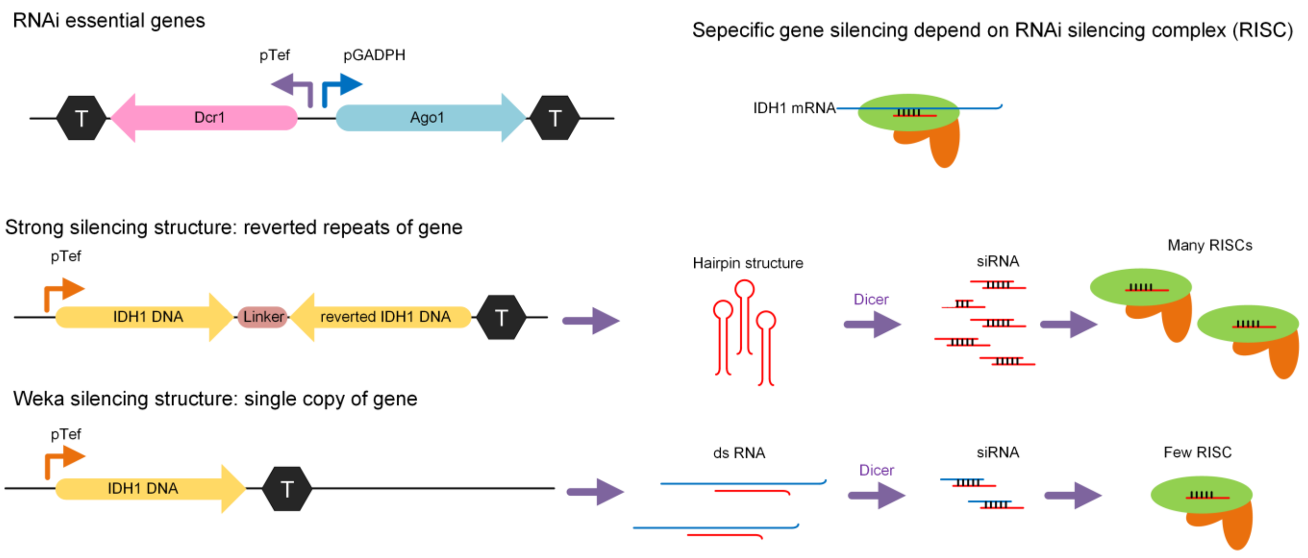


Fig. S5 RNAi was employed and established in this study.

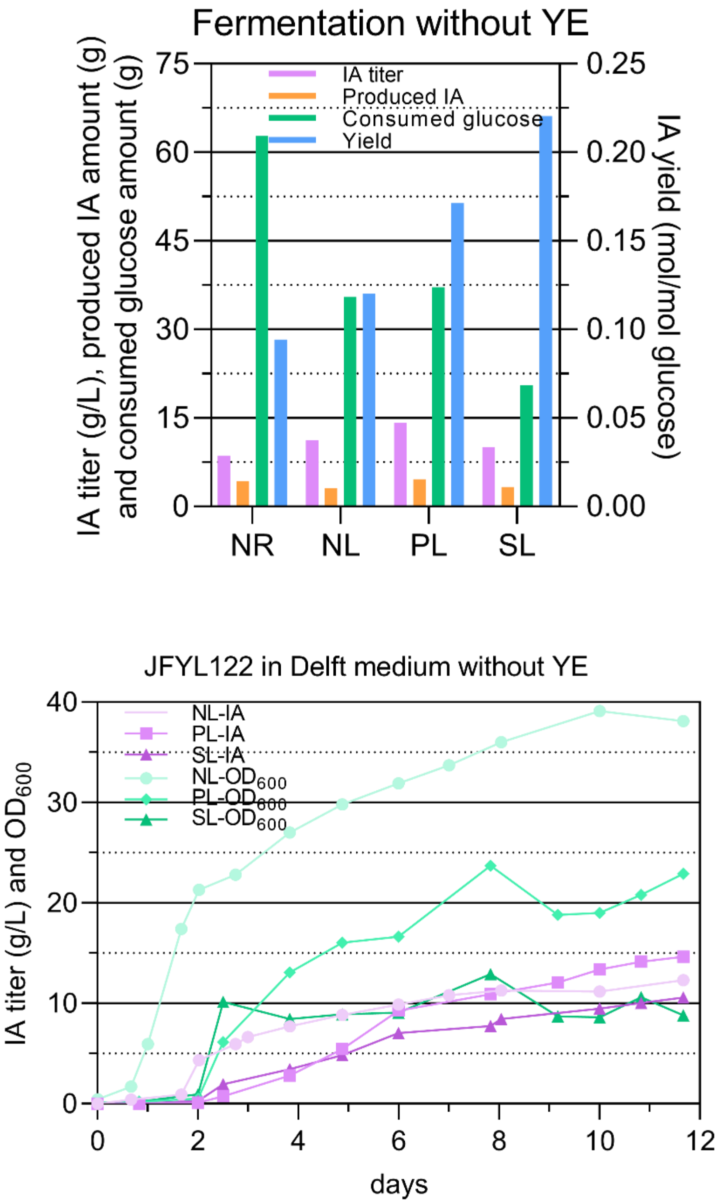


Fig. S6 JFYL122 was cultivated in 1L bioreactors without yeast extract. Top, itaconic acid production under nitrogen replete (C/N=22, $C_{100}N_{10}$), NL (C/N=88, $C_{100}N_{2.5}$), PL and SL conditions with pH at 5.5; Bottom, time course of itaconic acid titer and $OD_{600}$.

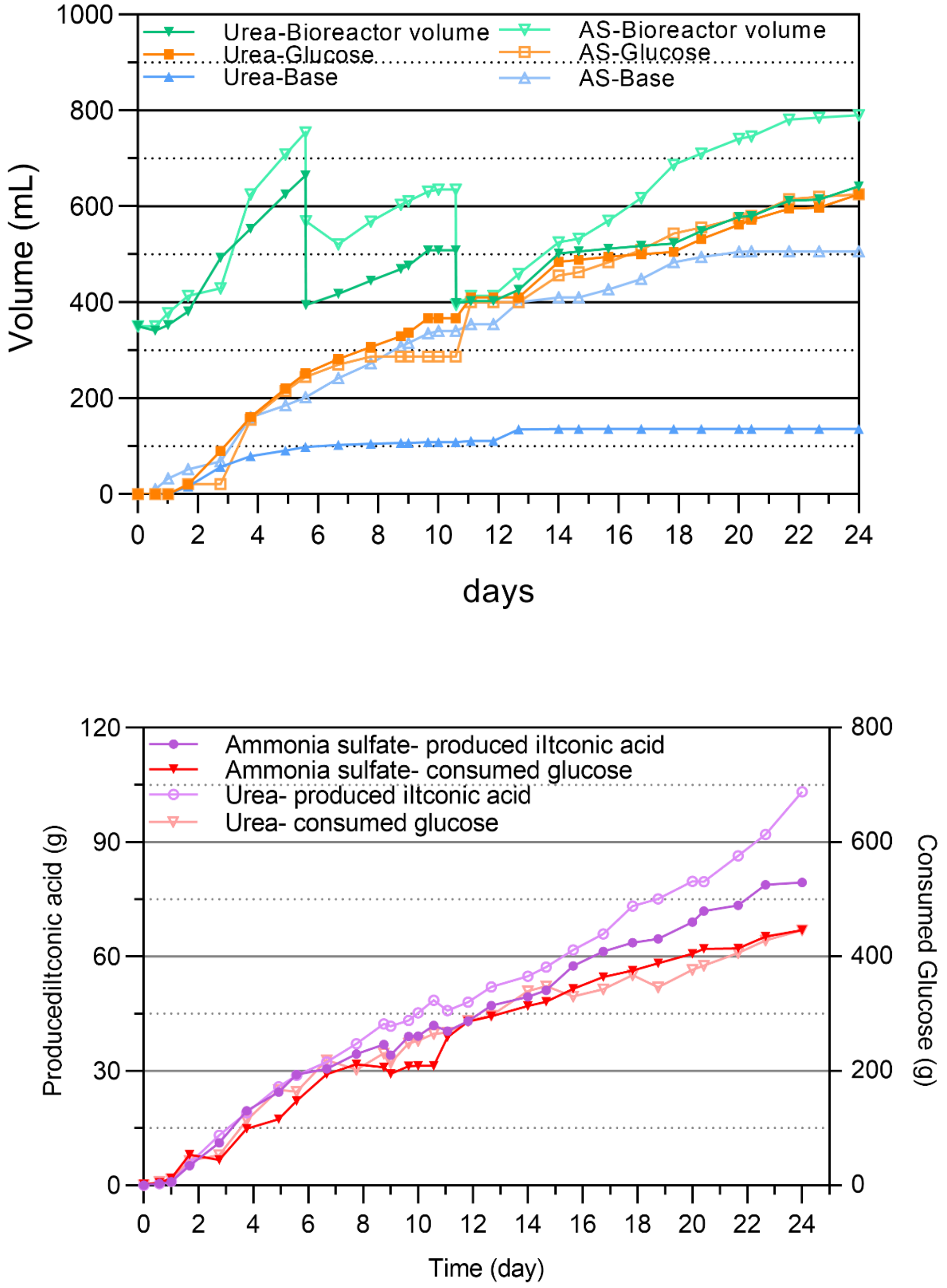


Fig. S7 produced IA and consumed glucose amount of JFYL121 when cultivated in 1L bioreactors with AS or urea. Top, volume of bioreactor, fed glucose and base; Bottom, Amount of produced itaconic acid and consumed glucose.

Table S1 Strains used in this study.

| Strains ID | Genotype | Reference |
|---|---|---|
| **WT** | W29 | [1] |
| **ST6512** | W29Δku70::cas9 | [1] |
| **OKYL029** | W29Δku70::cas9Δmhy1 | [2] |
| **JFYL007** | W29Δku70::cas9Δmhy1Δ*ARE1*Δ*LRO1*Δ*DGA1*Δ*DGA2* | This work |
| **JFYL008** | JFYL07, lntE3:p*BR30*-*AtCAD*-tlip2 | This work |
| **JFYL009** | JFYL07, lntE3:p*ICL1*-*AtCAD*-tlip2 | This work |
| **JFYL010** | JFYL07, lntE3:p4UASTef-*AtCAD*-tlip2 | This work |
| **JFYL013** | JFYL07, lntE3:pTef-*AtCAD*-tlip2 | This work |
| **JFYL014** | JFYL07, lntE1:pTef-*AtMTT*-tlip2, lntE3:pTef-*AtCAD*-tlip2 | This work |
| **JFYL016** | JFYL07Δ*ICL1* | This work |
| **JFYL017** | JFYL07Δ*ICL2* | This work |
| **JFYL018** | JFYL07Δ*ICL1*Δ*ICL2* | This work |
| **JFYL021** | JFYL07Δ*IDP* | This work |
| **JFYL023** | OKYL29, lntE3:pTef-*AtCAD*-tlip2 | This work |
| **JFYL025** | OKYL29, lntE1:pTef-*AtMTT*-tlip2, lntE3:pTef-*AtCAD*-tlip2 | This work |
| **JFYL028** | JFYL07Δ*ICL1*Δ*ICL2*Δ*IDP* | This work |
| **JFYL029** | JFYL07Δ*ICL1*, lntE3:pTef-*AtCAD*-tlip2 | This work |
| **JFYL030** | JFYL07Δ*ICL2*, lntE3:pTef-*AtCAD*-tlip2 | This work |
| **JFYL031** | JFYL07Δ*ICL1*Δ*ICL2*, lntE3:pTef-*AtCAD*-tlip2 | This work |
| **JFYL032** | JFYL07Δ*IDP*, lntE3:pTef-*AtCAD*-tlip2 | This work |
| **JFYL033** | JFYL07Δ*ICL1*Δ*ICL2*Δ*IDP*, lntE3:pTef-*AtCAD*-tlip2 | This work |
| **JFYL035** | JFYL07, lntE3:pTefin-*AtCAD*-tlip2 | This work |
| **JFYL036** | JFYL07, lntE1:pTefin-*AtMTT*-tlip2, lntE3:pTef-*AtCAD*-tlip2 | This work |
| **JFYL039** | JFYL07, lntE1:pTef-*AtMTT*-tlip2, lntC3:pTef-*AtCAD*-tlip2-p*GPD*-*AtMTT*-tpex20 | This work |
| **JFYL040** | JFYL07, lntE3:pTef-*AtCAD*-tlip2, lntC3:pTef-*AtCAD*-tlip2-p*GPD*-*AtMTT*-tpex20 | This work |
| **JFYL041** | JFYL07, lntE1:pTef-*AtMTT*-tlip2, lntE3:pTef-*AtCAD*-tlip2, lntC3:pTef-*AtCAD*-tlip2-p*GPD*-*AtMTT*-tpex20 | This work |
| **JFYL051** | JFYL07Δ*ICL1*Δ*ICL2*, lntC3:pTef-*AtCAD*-tlip2-p*GPD*-*AtMTT*-tpex20 | This work |
| **JFYL053** | JFYL07Δ*ICL1*Δ*ICL2*Δ*IDP*, lntC3:pTef-*AtCAD*-tlip2-p*GPD*-*AtMTT*-tpex20 | This work |
| **JFYL054** | JFYL07, lntE1:pTef-*AtMTT*-tlip2, lntE3:pTef-*AtCAD*-tlip2,lntC3:pTef-*AtACO*d-tlip2 | This work |
| **JFYL055** | JFYL07, lntE1:pTef-*AtMTT*-tlip2, lntE3:pTef-*AtCAD*-tlip2, lntC3:pTef-*AtACO*-tlip2 | This work |
| **JFYL056** | JFYL07, lntD1:pTef-Ylcox4-*AtCAD*-tlip2 | This work |
| **JFYL061** | JFYL07, lntE1:pTef-Akctp-tlip2, lntE3:pTef-*AtCAD*-tlip2 | This work |
| **JFYL062** | JFYL07, lntE1:pTefin-Akctp-tlip2, lntE3:pTef-*AtCAD*-tlip2 | This work |
| **JFYL063** | JFYL07, lntE1:pTef-Ummtt1-tlip2, lntE3:pTef-*AtCAD*-tlip2 | This work |
| **JFYL064** | JFYL07, lntE1:pTefin-Ummtt1-tlip2, lntE3:pTef-*AtCAD*-tlip2 | This work |
| **JFYL065** | JFYL07, lntC2:pTef-*AtCAD*-tlip2-p*GPD*-*AtMTT*-tpex20 | This work |
| **JFYL066** | JFYL07, lntC3:pTef-*AtCAD*-tlip2-p*GPD*-*AtMTT*-tpex20 | This work |
| **JFYL067** | JFYL07, lntD1:pTef-*AtCAD*-tlip2-p*GPD*-*AtMTT*-tpex20 | This work |
| **JFYL068** | JFYL07, lntE1:pTef-*AtCAD*-tlip2-p*GPD*-*AtMTT*-tpex20 | This work |
| **JFYL069** | JFYL07, lntE2:pTef-*AtCAD*-tlip2-p*GPD*-*AtMTT*-tpex20 | This work |
| **JFYL070** | JFYL07, lntE3:pTef-*AtCAD*-tlip2-p*GPD*-*AtMTT*-tpex20 | This work |
| **JFYL071** | JFYL07, lntC2:pTef-*AtCAD*-tlip2-p*GPD*-*AtMTT*-tpex20, lntC3:pTef-*AtCAD*-tlip2-p*GPD*-*AtMTT*-tpex20 | This work |
| **JFYL074** | JFYL07, lntC2:pTef-*AtCAD*-tlip2-p*GPD*-*AtMTT*-tpex20, lntC3:pTef-*AtCAD*-tlip2-p*GPD*-*AtMTT*-tpex20, lntD1:pTef-*AtCAD*-tlip2-p*GPD*-*AtMTT*-tpex20 | This work |
| **JFYL075** | JFYL07, lntC2:pTef-*AtCAD*-tlip2-p*GPD*-*AtMTT*-tpex20, lntC3:pTef-*AtCAD*-tlip2-p*GPD*-*AtMTT*-tpex20, lntD1:pTef-*AtCAD*-tlip2-p*GPD*-*AtMTT*-tpex20, lntE2:pTef-*AtCAD*-tlip2-p*GPD*-*AtMTT*-tpex20 | This work |

| | | |
|---|---|---|
| **JFYL076** | JFYL07, lntC2:pTef-*AtCAD*-tlip2-p*GPD*-*AtMTT*-tpex20, lntC3:pTef-*AtCAD*-tlip2-p*GPD*-*AtMTT*-tpex20, lntD1:pTef-*AtCAD*-tlip2-p*GPD*-*AtMTT*-tpex20, lntE1:pTef-*AtCAD*-tlip2-p*GPD*-*AtMTT*-tpex20, lntE1:pTef-*AtCAD*-tlip2-p*GPD*-*AtMTT*-tpex20 | This work |
| **JFYL077** | JFYL07, lntC2:pTef-*AtCAD*-tlip2-p*GPD*-*AtMTT*-tpex20, lntC3:pTef-*AtCAD*-tlip2-p*GPD*-*AtMTT*-tpex20, lntD1:pTef-*AtCAD*-tlip2-p*GPD*-*AtMTT*-tpex20, lntE1:pTef-*AtCAD*-tlip2-p*GPD*-*AtMTT*-tpex20, lntE2:pTef-*AtCAD*-tlip2-p*GPD*-*AtMTT*-tpex20, lntE3:pTef-*AtCAD*-tlip2-p*GPD*-*AtMTT*-tpex20 | This work |
| **JFYL083** | JFYL07, lntD1:pTef-Ylcox4-*AtCAD*-tlip2, lntE1:pTef-SceDIC-tlip2 | This work |
| **JFYL084** | JFYL07, lntD1:pTef-Ylcox4-*AtCAD*-tlip2, lntE1:pTefin-SceDIC-tlip2 | This work |
| **JFYL085** | JFYL07, lntD1:pTef-Ylcox4-*AtCAD*-tlip2, lntE1:pTef-MumSlc-tlip2 | This work |
| **JFYL086** | JFYL07, lntD1:pTef-Ylcox4-*AtCAD*-tlip2, lntE1:pTefin-MumSlc-tlip2 | This work |
| **JFYL087** | JFYL07, lntD1:pTef-Ylcox4-*AtCAD*-tlip2, lntE1:pTef-AthDtc-tlip2 | This work |
| **JFYL088** | JFYL07, lntD1:pTef-Ylcox4-*AtCAD*-tlip2, lntE1:pTefin-AthDtc-tlip2 | This work |
| **JFYL089** | JFYL07, lntD1:pTef-Ylcox4-*AtCAD*-tlip2, lntE3:pTef-*AtCAD*-tlip2, | This work |
| **JFYL090** | JFYL07, lntE3:pTef-UmTad1-tlip2 | This work |
| **JFYL091** | JFYL07, lntE1:pTef-*AtMTT*-tlip2, lntE3:pTef-UmTad1-tlip2 | This work |
| **JFYL092** | JFYL07, lntE1:pTef-UmTad1-tlip2, lntE3:pTef-UmTad1-tlip2 | This work |
| **JFYL093** | JFYL07, lntD1:pTef-*AtMTT*-tlip2, lntE1:pTef-UmTad1-tlip2, lntE3:pTef-UmTad1-tlip2 | This work |
| **JFYL094** | JFYL07, lntC3:pTef-*AtCAD*-tlip2-p*GPD*-*AtMTT*-tpex20, lntD1:pTef-YlAmpd-tlip2, | This work |
| **JFYL095** | JFYL07, lntC3:pTef-*AtCAD*-tlip2-p*GPD*-*AtMTT*-tpex20, p*IDH1*::p9 | This work |
| **JFYL096** | JFYL07, lntC3:pTef-*AtCAD*-tlip2-p*GPD*-*AtMTT*-tpex20, p*IDH1*::p10 | This work |
| **JFYL097** | JFYL07, lntD1:CRISPRi-*IDH1* | This work |
| **JFYL100** | JFYL07, lntC3:lip2t-*AtCAD*-Tefp-*GPD*p-*AtMTT*-pex20t, lntE1-pex2t-Dcr1-Tefp-*GPD*p-Ago1-lip2t | This work |
| **JFYL101** | JFYL07, lntC3:lip2t-*AtCAD*-Tefp-*GPD*p-*AtMTT*-pex20t, lntE1-pex2t-Dcr1-Tefp-*GPD*p-Ago1-lip2t, lntE3::Tefp-Ri1-pex20t | This work |
| **JFYL102** | JFYL07, lntC3:lip2t-*AtCAD*-Tefp-*GPD*p-*AtMTT*-pex20t, lntE1-pex2t-Dcr1-Tefp-*GPD*p-Ago1-lip2t, lntE3::Tefp-Ri2-pex20t | This work |
| **JFYL103** | JFYL07, lntC3:lip2t-*AtCAD*-Tefp-*GPD*p-*AtMTT*-pex20t, lntE1-pex2t-Dcr1-Tefp-*GPD*p-Ago1-lip2t, lntE3::Tefp-Ri3-pex20t | This work |
| **JFYL104** | JFYL07, lntC3:lip2t-*AtCAD*-Tefp-*GPD*p-*AtMTT*-pex20t, lntE1-pex2t-Dcr1-Tefp-*GPD*p-Ago1-lip2t, lntE3::Tefp-Ri4-pex20t | This work |
| **JFYL105** | JFYL07, lntC3:lip2t-*AtCAD*-Tefp-*GPD*p-*AtMTT*-pex20t, lntE1-pex2t-Dcr1-Tefp-*GPD*p-Ago1-lip2t, lntE3::Tefp-Ri5-pex20t | This work |
| **JFYL106** | JFYL07, lntC3:lip2t-*AtCAD*-Tefp-*GPD*p-*AtMTT*-pex20t, lntE1-pex2t-Dcr1-Tefp-*GPD*p-Ago1-lip2t, lntE3::Tefp-Ri6-pex20t | This work |
| **JFYL107** | JFYL07, lntC3:lip2t-*AtCAD*-Tefp-*GPD*p-*AtMTT*-pex20t, lntE1-pex2t-Dcr1-Tefp-*GPD*p-Ago1-lip2t, lntE3::Tefp-Ri7-pex20t | This work |
| **JFYL108** | JFYL07, lntC3:lip2t-*AtCAD*-Tefp-*GPD*p-*AtMTT*-pex20t, lntE1-pex2t-Dcr1-Tefp-*GPD*p-Ago1-lip2t, lntE3::Tefp-Ri8-pex20t | This work |
| **JFYL109** | JFYL07, lntC3:lip2t-*AtCAD*-Tefp-*GPD*p-*AtMTT*-pex20t, lntE1-pex2t-Dcr1-Tefp-*GPD*p-Ago1-lip2t, lntE3::Tefp-Ri9-pex20t | This work |
| **JFYL110** | JFYL07, lntC3:lip2t-*AtCAD*-Tefp-*GPD*p-*AtMTT*-pex20t, lntE1-pex2t-Dcr1-Tefp-*GPD*p-Ago1-lip2t, lntE3::Tefp-Ri10-pex20t | This work |
| **JFYL114** | JFYL07, lntE1:p4UASTef-*AtCAD*, lntE3:p4UASTef-*AtMTT* | This work |
| **JFYL115** | JFYL07, lntC3:pTef-*AtCAD*-tlip2-p*GPD*-*AtMTT*-tpex20, lntE1:p4UASTef-*AtCAD* | This work |
| **JFYL116** | JFYL07, lntC3:pTef-*AtCAD*-tlip2-p*GPD*-*AtMTT*-tpex20, lntE3:p4UASTef-*AtMTT* | This work |
| **JFYL117** | JFYL07, lntC2:pTef-*AtCAD*-tlip2-p*GPD*-*AtMTT*-tpex20, lntC3:pTef-*AtCAD*-tlip2-p*GPD*-*AtMTT*-tpex20, lntD1:pTef-*AtCAD*-tlip2-p*GPD*-*AtMTT*-tpex20, lntE2:pTef-*AtCAD*-tlip2-p*GPD*-*AtMTT*-tpex20, lntE1-pex2t-Dcr1-Tefp-*GPD*p-Ago1-lip2t, lntE3::Tefp-Ri3-pex20t | This work |
| **JFYL118** | JFYL07, lntC2:pTef-*AtCAD*-tlip2-p*GPD*-*AtMTT*-tpex20, lntC3:pTef-*AtCAD*-tlip2-p*GPD*-*AtMTT*-tpex20, lntD1:pTef-*AtCAD*-tlip2-p*GPD*-*AtMTT*-tpex20, lntE2:pTef-*AtCAD*-tlip2-p*GPD*-*AtMTT*-tpex20, lntE1-pex2t-Dcr1-Tefp-*GPD*p-Ago1-lip2t, lntE3::Tefp-Ri4-pex20t | This work |

| | | |
|---|---|---|
| **JFYL119** | JFYL07, lntC2:pTef-*AtCAD*-tlip2-p*GPD*-*AtMTT*-tpex20, lntC3:pTef-*AtCAD*-tlip2-p*GPD*-*AtMTT*-tpex20, lntD1:pTef-*AtCAD*-tlip2-p*GPD*-*AtMTT*-tpex20, lntE2:pTef-*AtCAD*-tlip2-p*GPD*-*AtMTT*-tpex20, p*IDH1*::p9 | This work |
| **JFYL120** | JFYL07, lntC2:pTef-*AtCAD*-tlip2-p*GPD*-*AtMTT*-tpex20, lntC3:pTef-*AtCAD*-tlip2-p*GPD*-*AtMTT*-tpex20, lntD1:pTef-*AtCAD*-tlip2-p*GPD*-*AtMTT*-tpex20, lntE2:pTef-*AtCAD*-tlip2-p*GPD*-*AtMTT*-tpex20, p*IDH1*::p10 | This work |
| **JFYL121** | JFYL07, lntC2:pTef-*AtCAD*-tlip2-p*GPD*-*AtMTT*-tpex20, lntC3:pTef-*AtCAD*-tlip2-p*GPD*-*AtMTT*-tpex20, lntD1:pTef-*AtCAD*-tlip2-p*GPD*-*AtMTT*-tpex20, lntE2:pTef-*AtCAD*-tlip2-p*GPD*-*AtMTT*-tpex20, lntE1-p4UASTef-*AtMTT*-pex2t | This work |
| **JFYL122** | JFYL07, lntC2:pTef-*AtCAD*-tlip2-p*GPD*-*AtMTT*-tpex20, lntC3:pTef-*AtCAD*-tlip2-p*GPD*-*AtMTT*-tpex20, lntD1:pTef-*AtCAD*-tlip2-p*GPD*-*AtMTT*-tpex20, lntE1:pTef-*AtCAD*-tlip2-p*GPD*-*AtMTT*-tpex20, lntE2:pTef-*AtCAD*-tlip2-p*GPD*-*AtMTT*-tpex20, lntE3:pTef-*AtCAD*-tlip2-p*GPD*-*AtMTT*-tpex20, p*IDH1*::p9 | This work |
| **JFYL123** | JFYL07, lntC2:pTef-*AtCAD*-tlip2-p*GPD*-*AtMTT*-tpex20, lntC3:pTef-*AtCAD*-tlip2-p*GPD*-*AtMTT*-tpex20, lntD1:pTef-*AtCAD*-tlip2-p*GPD*-*AtMTT*-tpex20, lntE1:pTef-*AtCAD*-tlip2-p*GPD*-*AtMTT*-tpex20, lntE2:pTef-*AtCAD*-tlip2-p*GPD*-*AtMTT*-tpex20, lntE3:pTef-*AtCAD*-tlip2-p*GPD*-*AtMTT*-tpex20, p*IDH1*::p10 | This work |
| **JFYL124** | JFYL07, lntE1-pex2t-Dcr1-Tefp-*GPD*p-Ago1-lip2t, lntE3::Tefp-Ri3-pex20t | This work |
| **JFYL125** | JFYL07, lntE1-pex2t-Dcr1-Tefp-*GPD*p-Ago1-lip2t, lntE3::Tefp-Ri4-pex20t | This work |
| **JFYL128** | JFYL07, lntE1:3*pTef-*AtCAD*-GSG-*AtACO*d-tlip2, lntD1:3*pTef-*AtCAD*-GSG-*AtACO*d-tlip2 | This work |
| **JFYL129** | JFYL07, lntE1:3*pTef-*AtCAD*-GSG-*AtACO*d-tlip2, lntD1:3*pTef-*AtCAD*-GSG-*AtACO*d-tlip2, b lntC3:pTef-*AtACO*d-tlip2 | This work |
| **JFYL130** | JFYL07, lntD1:pTef-*AtACO*d-GSG-*AtCAD*-tlip2 | This work |
| **JFYL131** | JFYL07, lntD1:pTef-cox4-*AtCAD*-GSG-*AtACO*d-tlip2 | This work |
| **JFYL132** | JFYL07, lntD1:pTef-*AtACO*-GSG-*AtCAD*-tlip2 | This work |
| **JFYL133** | JFYL07, lntE1:pTef-*AtMTT*, lntD1:pTef-*AtCAD*-GSG-*AtACO*d-tlip2 | This work |
| **JFYL134** | JFYL07, lntE1:pTef-*AtMTT*, lntD1:pTef-*AtACO*d-GSG-*AtCAD*-tlip2 | This work |
| **JFYL135** | JFYL07, lntE1:pTef-*AtMTT*, lntD1:pTef-cox4-*AtCAD*-GSG-*AtACO*d-tlip2 | This work |
| **JFYL136** | JFYL07, lntE1:pTef-*AtMTT*, lntD1:pTef-*AtACO*-GSG-*AtCAD*-tlip2 | This work |

Table S2 Plasmids used in this study.

| Plasmid ID | Full name | Purpose/comments | Reference |
|---|---|---|---|
| OKEC003 | pgRNA-*ARE1* | *ARE1* deletion | [3] |
| JFEC001 | pgRNA-*LRO1* | *LRO1* deletion | This study |
| JFEC002 | pgRNA-*DGA1* | *DGA1* deletion | This study |
| JFEC003 | pgRNA-*DGA2* | *DGA2* deletion | This study |
| JFEC004 | pgRNA-*ICL1* | *ICL1* deletion | This study |
| JFEC005 | pgRNA-*ICL2* | *ICL2* deletion | This study |
| JFEC006 | pgRNA-*IDH1* | *IDH1* deletion | This study |
| JFEC007 | pgRNA-*IDH2* | *IDH2* deletion | This study |
| JFEC008 | pgRNA-*IDP* | *IDP* deletion | This study |
| JFEC011 | pE3-p*BR30*-*AtCAD* | Overexpression of *AtCAD* under *BR30* promoter | This study |
| JFEC012 | pE3-p*ICL2*-*AtCAD* | Overexpression of *AtCAD* under *ICL2* promoter | This study |
| JFEC013 | pE3-pTef-*AtCAD* | Overexpression of *AtCAD* under Tef promoter | This study |
| JFEC014 | pE3-p4UASTef-*AtCAD* | Overexpression of *AtCAD* under 4UASTef promoter | This study |
| JFEC015 | pE3-pTefin-*AtCAD* | Overexpression of *AtCAD* under *BR30* promoter | This study |
| JFEC017 | pD1-pTef-cox4-*AtCAD* | Overexpression of *AtCAD* with cox4 mitochondrial leading sequence under Tef promoter | This study |
| JFEC021 | pE1-pTef-*AtMTT* | Overexpression of *AtMTT* under Tef promoter | This study |
| JFEC022 | pE1-p4UASTef-*AtMTT* | Overexpression of *AtMTT* under 4UASTef promoter | This study |
| JFEC023 | pE1-pTefin-*AtMTT* | Overexpression of *AtMTT* under Tef intron promoter | This study |
| JFEC024 | pC2-pTef-*AtCAD*-tlip2-p*GPD*-*AtMTT*-tpex20 | Overexpression of *AtCAD* under Tef promoter and *AtMTT* under *GPD* promoter at lntC2 locus | This study |
| JFEC025 | pC3-pTef-*AtCAD*-tlip2-p*GPD*-*AtMTT*-tpex20 | Overexpression of *AtCAD* under Tef promoter and *AtMTT* under *GPD* promoter at lntC3 locus | This study |
| JFEC026 | pD1-pTef-*AtCAD*-tlip2-p*GPD*-*AtMTT*-tpex20 | Overexpression of *AtCAD* under Tef promoter and *AtMTT* under *GPD* promoter at lntD1 locus | This study |
| JFEC027 | pE1-pTef-*AtCAD*-tlip2-p*GPD*-*AtMTT*-tpex20 | Overexpression of *AtCAD* under Tef promoter and *AtMTT* under *GPD* promoter at lntE1 locus | This study |
| JFEC028 | pE2-pTef-*AtCAD*-tlip2-p*GPD*-*AtMTT*-tpex20 | Overexpression of *AtCAD* under Tef promoter and *AtMTT* under *GPD* promoter at lntE2 locus | This study |
| JFEC029 | pE3-pTef-*AtCAD*-tlip2-p*GPD*-*AtMTT*-tpex20 | Overexpression of *AtCAD* under Tef promoter and *AtMTT* under *GPD* promoter at lntE3 locus | This study |
| JFEC031 | pC3-pTef-MuSlc | Overexpression of *MuSlc* under Tef promoter | This study |
| JFEC032 | pC3-pTefin-MuSlc | Overexpression of *MuSlc* under Tef intron promoter | This study |
| JFEC033 | pC3-pTef-AthDlc | Overexpression of *AthDlc* under Tef promoter | This study |
| JFEC034 | pC3-pTefin-AthDlc | Overexpression of *AthDlc* under Tef intron promoter | This study |
| JFEC035 | pC3-pTef-SceDlc | Overexpression of *SceDlc* under Tef promoter | This study |
| JFEC036 | pC3-pTefin-SceDlc | Overexpression of *SceDlc* under Tef intron promoter | This study |
| JFEC037 | pE1-pTef-Ummtt1 | Overexpression of *Ummtt1* under Tef promoter | This study |
| JFEC038 | pE1-pTefin-Ummtt1 | Overexpression of *Ummtt1* under Tef intron promoter | This study |
| JFEC039 | pE1-pTef-AkCtp | Overexpression of *MuSlc* under Tef promoter | This study |
| JFEC040 | pE1-pTefin-AkCtp | Overexpression of *MuSlc* under Tef intron promoter | This s study |
| JFEC041 | pC3-pTef-*AtACO*d | Overexpression of *AtACO*d under Tef promoter | This study |
| JFEC042 | pC3-pTef-*AtACO* | Overexpression of *AtACO* under Tef promoter | This study |
| JFEC043 | pD1-pTef-*AtACO*d-GSG-*AtCAD* | Overexpression of *AtACO*d-*AtCAD* fused protein with a GSG linker under Tef promoter | This study |
| JFEC044 | pD1-pTef-*AtCAD*-GSG-*AtACO*d | Overexpression of *AtCAD*-*AtACO*d fused protein with a GSG linker under Tef promoter | This study |
| JFEC045 | pD1-pTef-cox4-GSG-*AtACO*d | Overexpression of *AtCAD*-*AtACO*d fused protein with a GSG linker and with cox4 mitochondrial leading sequence under Tef promoter | This study |

| | | | |
|---|---|---|---|
| JFEC046 | pD1-pTef-*AtACO*-GSG-*AtCAD* | Overexpression of *AtACO*-*AtCAD* fused protein with a GSG linker under Tef promoter | This study |
| JFEC100 | plntE1-Pex20t-Dcr1-Tefp-*GPD*p-Ago1-lip2t | For RNAi establishment, overexpression of Dcr1 under Tef promoter and Ago1 under *GPD* promoter | This study |
| JFEC101 | plntE3-Tefp-*IDH1*-inverted repeats1 | Overexpression of *IDH1* inverted repeats 1, which is 170bp | This study |
| JFEC102 | plntE3-Tefp-*IDH1*-inverted repeats2 | Overexpression of *IDH1* inverted repeats 2, which is 500bp | This study |
| JFEC103 | plntE3-Tefp-*IDH1*-inverted repeats3 | Overexpression of *IDH1* inverted repeats 3, which is 1k | This study |
| JFEC104 | plntE3-Tefp-*IDH1*-inverted repeats4 | Overexpression of *IDH1* inverted repeats 4, which is 1742bp (full length) | This study |
| JFEC105 | plntE3-Tefp-*IDH1*-inverted repeats5 | Overexpression of *IDH1* inverted repeats 5, which is 170bp | This study |
| JFEC106 | plntE3-Tefp-*IDH1*-inverted repeats6 | Overexpression of *IDH1* inverted repeats 6, which is 170bp | This study |
| JFEC107 | plntE3-Tefp-*IDH1*-inverted repeats7 | Overexpression of *IDH1* inverted repeats 7, which is | This study |
| JFEC108 | plntE3-Tefp-*IDH1*-single1 | Overexpression of *IDH1* single repeats 1, which is 170bp from N terminal | This study |
| JFEC109 | plntE3-Tefp-*IDH1*-single2 | Overexpression of *IDH1* single repeats 2, which is 1742bp full length of cDNA | This study |
| JFEC110 | plntE3-Tefp-*IDH1*-single3 | Overexpression of *IDH1* single repeats 3, which is 170bp from C terminal | This study |
| JFEC113 | plntE1-Tefp-Adi1-lip2t | Overexpression of Adi1 under Tef promoter | This study |
| JFEC114 | plntE3-Tefp-Tad1-lip2t | Overexpression of Tad1 under Tef promoter | This study |
| JFEC115 | plntE1-4UASTefp-*AtMTT*-lip2t | Overexpression of *AtMTT* under 4UASTefp promoter | This study |
| JFEC116 | plntE3-4UASTefp-*AtCAD*-lip2t | Overexpression of *AtCAD* under 4UASTefp promoter | This study |
| JFEC117 | pD1-pTef-AMPD | Overexpression of native AMPD under Tef promoter | This study |
| JFEC127 | pD1-3*(Pex20t-*AtCAD*-Tefp-*GPD*p-*AtMTT*-lip2t) | Overexpression of 3 cassettes at lntD1 locus, and each cassette consists *AtCAD* under Tef promoter and *AtMTT* under *GPD* promoter | This study |
| JFEC158 | pCRISPRi ddcas12a | For CRISPRi establishment, overexpression of ddcas12a and 4 gRNAs for *IDH1* b | This study |
| JFEC159 | pE1-Pex20t-*AtCAD*-Tefp-*GPD*p-*AtMTT*-lip2t*3 | Overexpression of 3 cassettes at lntE1 locus, and each cassette consists *AtCAD* under Tef promoter and *AtMTT* under *GPD* promoter | This study |

Table S3 Primers used in this study.

| Primer ID | Oligos | Comments |
|---|---|---|
| F001=*LRO1*-Del-cr-F | ACTCAACGCCAAGTACCCGGgttttagagct | gRNA expression cassette for *DGA1* deletion for *LRO1* deletion with 20 bp site-specific gRNA BioBrick |
| F002=*LRO1*-Del-cr-R | CCGGGTACTTGGCGTTGAGTtaaccaacct | For *LRO1* gRNA plasmids with 20 bp site-specific gRNA BioBrick |
| F007=*LRO1*-Del-ck-F | CTGAATTTCCCGATTTATTC | check primer for *LRO1* deletion |
| F008=*LRO1*-Del-ck-R | GAAACGCGCATATGATAGTG | check primer for *LRO1* deletion |
| F009=*DGA1*-Del-ck-F | CGAGCGAATCGCACACAAAC | check primer for *DGA1* deletion |
| F010=*DGA1*-Del-ck-R | CCATGTATGACATTCGAGCC | check primer for *DGA1* deletion |
| F011=*DGA2*-Del-ck-F | CAAACGAGTATACTTGTAGC | check primer for *DGA2* deletion |
| F012=*DGA2*-Del-ck-R | CACAGTCACGAAAACCATAC | check primer for *DGA2* deletion |
| F013=gRNA-cass1-F | cgtgcgaUagtgaatcattgctaacagatc | PR-10607 in EasyClone |
| F014=gRNA-cass1-R | cacgcgaUaccgtacccacacaaaaaaagcaccaccgactc | PR-10604 in EasyClone |
| F017=gRNA-cass2-F | AGTGCAGGUagtgaatcattgctaacagatc | PR-15790 in EasyClone |
| F018=gRNA-cass2-R | ACCTGCACUaccgtacccacacaaaaaaagcac | PR-15791 in EasyClone |
| F027=*LRO1*-RM-F | TAAAAAAAGTGTAATCGGCTTTTTCCGGTTGATCACAACCATCAAGAGTCCGTTTTGTAGAGTAATATGTTTTGTATATCACACTGATG | Repair fragments for *LRO1* deletion |
| F028=*LRO1*-RM-R | CATCAGTGTGATATACAAAACATATTACTCTACAAAACGGACTCTTGATGGTTGTGATCAACCGGAAAAAGCCGATTACACTTTTTTA | Repair fragments for *LRO1* deletion |
| F029=*DGA1*-RM-F | CAAAAAAATACTCATTAGCTATTTGCCTAACCCAGGCAGTTTTCCAGCTTTTGTTTTGTGTGACTTGTCTGTTGCCTGTTGTTAGAAGAA | Repair fragments for *DGA1* deletion |
| F030=*DGA1*-RM-R | TTCTTCTAACAACAGGCAACAGACAAGTCACACAAAACAAAAGCTGGAAAACTGCCTGGGTTAGGCAAATAGCTAATGAGTATTTTTTTG | Repair fragments for *DGA1* deletion |
| F031=*DGA2*-RM-F | TGTCAATATTATTTATCACTGTAAAGGCTACTGATGAGTGTTATGTTTGCGGGCGGTACGGGTACAGCGACTTTGGGTGTAGCTATGGTG | Repair fragments for *DGA2* deletion |
| F032=*DGA2*-RM-R | CACCATAGCTACACCCAAAGTCGCTGTACCCGTACCGCCCGCAAACATAACACTCATCAGTAGCCTTTACAGTGATAAATAATATTGACA | Repair fragments for *DGA2* deletion |
| F033=gRNA-cass-ck-F | TTGGAGGCGACGTGGCAG | Check primer for gRNA plasmids |
| F034=gRNA-cass-ck-R | AAATGCGGCCGCGAATGC | Check primer for gRNA plasmids |
| F035=*DGA1*-Del-cr2-F | GGCGTCTTCAACTACGATGTgttttagagct | gRNA expression cassette for *DGA1* deletion |
| F036=*DGA1*-Del-cr2-R | ACATCGTAGTTGAAGACGCCtaaccaacct | gRNA expression cassette for *DGA1* deletion |
| F037=*DGA1*-Del-cr3-F | AGCATAGCATGAAAATTGTGgttttagagct | gRNA expression cassette for *DGA1* deletion |
| F038=*DGA1*-Del-cr3-R | CACAATTTTCATGCTATGCTtaaccaacct | gRNA expression cassette for *DGA1* deletion |
| F039=*DGA2*-Del-cr2-F | GAGCCAGACCATCATAGACAgttttagagct | gRNA expression cassette for *DGA2* deletion |
| F040=*DGA2*-Del-cr2-R | TGTCTATGATGGTCTGGCTCtaaccaacct | gRNA expression cassette for *DGA2* deletion |
| F041=*DGA2*-Del-cr3-F | GAGCCAGACCATCATAGACAgttttagagct | gRNA expression cassette for *DGA2* deletion |
| F042=*DGA2*-Del-cr3-R | TGTCTATGATGGTCTGGCTCtaaccaacct | gRNA expression cassette for *DGA2* deletion |
| F049=ICD1-RM-F | TTTTTTTCTTGTCGCCAACTTCGTGGAACCCCCAAAGAAATCACAACGATATAACGATAATGATAATGATTTGATGTAATGTTGGAAGCG | Repair fragments |
| F050=ICD1-RM-R | CGCTTCCAACATTACATCAAATCATTATCATTATCGTTATATCGTTGTGATTTCTTTGGGGGTTCCACGAAGTTGGCGACAAGAAAAAAA | Repair fragments |
| F051=ICD2-RM-F | TCAATAATTCGTTCAATGTCATAATATCATCTAAATCATACATAATGTGGATGTGGATATGTTTTCGATTAGACGACAAGATGTTGCCAA | Repair fragments |

| | | |
|---|---|---|
| F052=ICD2-RM-R | TTGGCAACATCTTGTCGTCTAATCGAAAACATATCCACATCCACATTATGTATGATTTAGATGATATTATGACATTGAACGAATTATTGA | Repair fragments |
| F053=ICD3-RM-F | ATCGTGCGGTTGTGTCGCATAAATCATTAATATAAACATTTTCCCGGCTGGCAAGGGAGGGTCTGTGGGGGTTCAACGTGGGGTTGCATG | Repair fragments |
| F054=ICD3-RM-R | CATGCAACCCCACGTTGAACCCCCACAGACCCTCCCTTGCCAGCCGGGAAAATGTTTATATTAATGATTTATGCGACACAACCGCACGAT | Repair fragments |
| F055=*ICL1*-RM-F | GCCTCCTTCACCTACGGTGCCCTCGACCCCGTCCAGGTGACCCAGGCAGTTTGTTTAGCAAAATATATTTAACGAGTTTGATAGAGGCGC | Repair fragments |
| F056=*ICL1*-RM-R | GCGCCTCTATCAAACTCGTTAAATATATTTTGCTAAACAAACTGCCTGGGTCACCTGGACGGGGTCGAGGGCACCGTAGGTGAAGGAGGC | Repair fragments |
| F057=*ICL2*-RM-F | AACACGAGCGGTTGCTGATTCGGTCACTAAAAAACAAGTGCAAAAGCATGGTACGATGTGTAACTAGCTAACTGGATGAAAGAACGGCAA | Repair fragments |
| F058=*ICL2*-RM-R | TTGCCGTTCTTTCATCCAGTTAGCTAGTTACACATCGTACCATGCTTTTGCACTTGTTTTTTAGTGACCGAATCAGCAACCGCTCGTGTT | Repair fragments |
| F059=ICD1-Del-cr1-F | GGGTCGCACAGCTCGAAGGGgttttagagct | gRNA expression cassette for deletion |
| F060=ICD1-Del-cr1-R | CCCTTCGAGCTGTGCGACCCtaaccaacct | gRNA expression cassette for deletion |
| F061=ICD1-Del-cr2-F | GTGTCGCAGCATCATCACGGgttttagagct | gRNA expression cassette for deletion |
| F062=ICD1-Del-cr2-R | CCGTGATGATGCTGCGACACtaaccaacct | gRNA expression cassette for deletion |
| F063=ICD2-Del-cr1-F | ACTCACCTTAGCCAGGTAGGgttttagagct | gRNA expression cassette for deletion |
| F064=ICD2-Del-cr1-R | CCTACCTGGCTAAGGTGAGTtaaccaacct | gRNA expression cassette for deletion |
| F065=ICD2-Del-cr2-F | GAGGTGTCGCAGAAGCATGGgttttagagct | gRNA expression cassette for deletion |
| F066=ICD2-Del-cr2-R | CCATGCTTCTGCGACACCTCtaaccaacct | gRNA expression cassette for deletion |
| F067=ICD3-Del-cr1-F | TGGTGGACATGGTTTTAGAAgttttagagct | gRNA expression cassette for deletion |
| F068=ICD3-Del-cr1-R | TTCTAAAACCATGTCCACCAtaaccaacct | gRNA expression cassette for deletion |
| F069=ICD3-Del-cr2-F | ATCCTCGTATGTGCTGACCGgttttagagct | gRNA expression cassette for deletion |
| F070=ICD3-Del-cr2-R | CGGTCAGCACATACGAGGATtaaccaacct | gRNA expression cassette for deletion |
| F071=*ICL1*-Del-cr1-F | CGAGGCATGAAGGCTTACGGgttttagagct | gRNA expression cassette for deletion |
| F072=*ICL1*-Del-cr1-R | CCGTAAGCCTTCATGCCTCGtaaccaacct | gRNA expression cassette for deletion |
| F073=*ICL1*-Del-cr2-F | GCTCCCGGTACCAAGAAGTGgttttagagct | gRNA expression cassette for deletion |
| F074=*ICL1*-Del-cr2-R | CACTTCTTGGTACCGGGAGCtaaccaacct | gRNA expression cassette for deletion |
| F075=*ICL2*-Del-cr1-F | CGAGATGATGACCTACGACGgttttagagct | gRNA expression cassette for deletion |
| F076=*ICL2*-Del-cr1-R | CGTCGTAGGTCATCATCTCGtaaccaacct | gRNA expression cassette for deletion |
| F077=*ICL2*-Del-cr2-F | GTGGATGTCTCTGTCCCACGgttttagagct | gRNA expression cassette for deletion |
| F078=*ICL2*-Del-cr2-R | CGTGGGACAGAGACATCCACtaaccaacct | gRNA expression cassette for deletion |
| F079=ICD1-Del-ck-F | AGGGGAGTTAGACGGACGTGGG | gRNA expression cassette for deletion |
| F080=ICD1-Del-ck-R | CTTCCGCACATCCTTTCCACCC | gRNA expression cassette for deletion |
| F081=ICD2-Del-ck-F | TACCCAACTCCATCACACCCAG | gRNA expression cassette for deletion |
| F082=ICD2-Del-ck-R | GGTCGGGTACCCGGACGGATGT | gRNA expression cassette for deletion |
| F083=ICD3-Del-ck-F | GAGGTGGACTTCCCTGTGCGAG | gRNA expression cassette for deletion |
| F084=ICD3-Del-ck-R | GCATAATCTGTGCTGCTTCCCC | gRNA expression cassette for deletion |
| F085=*ICL1*-Del-ck-F | GGATGAGGTGTTGTGTGGTGGG | gRNA expression cassette for deletion |
| F086=*ICL1*-Del-ck-R | CACCTCACCTCCCTCACCCCTT | gRNA expression cassette for deletion |
| F087=*ICL2*-Del-ck-F | GGAGCAAACTGCAGGACGGATG | gRNA expression cassette for deletion |
| F088=*ICL2*-Del-ck-R | CGTACGATCACGATGTGTGGGT | gRNA expression cassette for deletion |
| F097=ACOdL1CAD-C3tef-PF | TTCAACGGAATGCGTGCGATAGAGACCGGGTTGGCGGCGC | *AtACO* and *AtCAD* fused protein |
| F098=ACOdL1CAD-C3tef-G1R | TGCTTGGTCATACCAGAACCGTTGGAGGCGGCCTTTCGAG | *AtACO* and *AtCAD* fused protein |
| F099=ACOdL1CAD-C3tef-G2F | CTCGAAAGGCCGCCTCCAACGGTTCTGGTATGACCAAGCAGTCTGCCGAC | *AtACO* and *AtCAD* fused protein |

| | | |
|---|---|---|
| F100=CADL1CAOd-D1tef-PF | TTCAACGGAATGCGTGCGATCGCTTGAGGATCCAGAGACCGGGTTGGCGGCGC | *AtACO* and *AtCAD* fused protein |
| F101=CADL1ACOd-D1tef-G1R | GCCACGGTAGCACCAGAACCCACCAGGGGAGACTTCACGG | *AtACO* and *AtCAD* fused protein |
| F102=CADL1ACOd-D1tef-G2F | CCGTGAAGTCTCCCCTGGTGGGTTCTGGTGCTACCGTGGCCGACTCGCC | *AtACO* and *AtCAD* fused protein |
| F106=*AtCAD*-USER-PF | CGTGCGAUAGAGACCGGGTTGGCGGCGC | Promoter amplification |
| F107=*AtCAD*-USER-PR | ATGACAGAUTTTGAATGATTCTTATACTC | Promoter amplification |
| F108=*AtCAD*-USER-GF | ATCTGTCAUGCCACAATGACCAAGCAGTCTGCCGACTC | Gene amplification |
| F109=*AtCAD*-USER-GR | CACGCGAUTTACACCAGGGGAGACTTCACG | Gene amplification |
| F115=Tefin-PR | CTGCGGTTAGTACTGCAAAAAGTGCTGGTC | Tef intron promoter amplification |
| F120=PR-14617-ckF | tatccctgtgttgaatc | Check primer for gene integration |
| F121=PR-14619-ckR | tatcgacccagttagc | Check primer for gene integration |
| F128=*AtCAD*-g2-tef-cox4-PR | TTGAAGGCGAGCATTGTGGCTTTGAATGATTCTTATACTC | Cox4-*AtCAD* promoter amplification |
| F129=*AtCAD*-g2-tef-cox4-GF | GCCACAATGCTCGCCTTCAAGTCTCTCCGACCCTCTGCTGTCTCCCGACTGGCAACCTCCACCCGAGCTGCCCACGTCATCTCCACCAAGCAGTCTGCCGACTC | Cox4-*AtCAD* gene amplification |
| F148=INT-C2-ck-F | AGACGCGAAGGACGACATCC | Check primer for intergration |
| F149=INT-C2-ck-R | TCGGCATCTTCATTCAGCTG | Check primer for intergration |
| F150=INT-C3-ck-F | CTCTCCACATTTCCAGATAG | Check primer for intergration |
| F151=INT-C3-ck-R | CTATCTAATTTCGTGGCCGC | Check primer for intergration |
| F152=INT-D1-ck-F | GTCGGAAGATGAACGCAATC | Check primer for intergration |
| F153=INT-D1-ck-R | ATGGCACATGTGCTAGAATG | Check primer for intergration |
| F154=INT-E1-ck-F | TACACAGGCTCTTCACTCAC | Check primer for intergration |
| F155=INT-E1-ck-R | CTGGAGATTCTCTCCTAGCT | Check primer for intergration |
| F156=INT-E2-ck-F | CAACTCCGTCTGGTGTCTCC | Check primer for intergration |
| F157=INT-E2-ck-R | CGGTCACCATCGCGGTTCTC | Check primer for intergration |
| F158=INT-E3-ck-F | ATGGGCCACATTGTGACACC | Check primer for intergration |
| F159=INT-E3-ck-R | CAATCTGGGGAACTTCGCGT | Check primer for intergration |
| F162=Del-ICD1-RM-UF | ttgcggcacaatccggatat | 1000 bp repair fragment of deletion |
| F163=Del-ICD1-RM-UR | CATTATCATTATCGTTATATCGTtgtgatttctttgggggttc | 1000 bp repair fragment of deletion |
| F164=Del-ICD1-RM-LF | GAACCCCCAAAGAAATCACAacgatataacgataatgataatg | 1000 bp repair fragment of deletion |
| F165=Del-ICD1-RM-LR | ttagcgtcaactggaaggcg | 1000 bp repair fragment of deletion |
| F166=Del-ICD1-RM-FsnF | ggcactgtgttctatttctg | 1000 bp repair fragment of deletion |
| F167=Del-ICD1-RM-FsnR | caacaagttagttgggtggg | 1000 bp repair fragment of deletion |
| F168=Del-ICD2-RM-UF | ttgggtgactgatgtgtgtg | 1000 bp repair fragment of deletion |
| F169=Del-ICD2-RM-UR | CGAAAACATATCCACATCCACAttatgtatgatttagatgatattatgac | 1000 bp repair fragment of deletion |
| F170=Del-ICD2-RM-LF | GTCATAATATCATCTAAATCATACATAAtgtggatgtggatatgttttcg | 1000 bp repair fragment of deletion |
| F171=Del-ICD2-RM-LR | ttctgccctgatatcgcgag | 1000 bp repair fragment of deletion |
| F172=Del-ICD2-RM-FsnF | aaagggtatcacgtgtcgag | 1000 bp repair fragment of deletion |
| F173=Del-ICD2-RM-FsnR | ggattctgtgtgcgcgttcg | 1000 bp repair fragment of deletion |
| F196=ICD1-RM-F1 | CTCGTCGACATCACTTTTTTTTCTTGTCGCCAACTTCGTGGAACCCCCAAAGAAATCACAACGATATAACGATAATGATAATGATTTGATGTAATGTTGGAAGCGATATGTGTGATCATC | Repair fragment for deletion |

| | | |
|---|---|---|
| F197=ICD1-RM-R1 | GATGATCACACATATCGCTTCCAACATTACATCAAATCATTATC ATTATCGTTATATCGTTGTGATTTCTTTGGGGGTTCCACGAAGTT GGCGACAAGAAAAAAAAGTGATGTCGACGAG | Repair fragment for deletion |
| F198=ICD2-RM-F1 | CACACCCACTACTCATCAATAATTCGTTCAATGTCATAATATCAT CTAAATCATACATAATGTGGATGTGGATATGTTTTCGATTAGAC GACAAGATGTTGCCAAAAGGTGTCCTAGTTA | Repair fragment for deletion |
| F199=ICD2-RM-R1 | TAACTAGGACACCTTTTGGCAACATCTTGTCGTCTAATCGAAAA CATATCCACATCCACATTATGTATGATTTAGATGATATTATGACA TTGAACGAATTATTGATGAGTAGTGGGTGTG | Repair fragment for deletion |
| F200=*ICL1*-RM-F1 | CAAAGAAGTCGGTCTCACCAATGCAAGTGTCACATCAAACATCT GTCCCGTACTAACCCAGCAGTTTGTTTAGCAAAATATATTTAAC GAGTTTGATAGAGGCGCTGGACTACATAATTA | Repair fragment for deletion |
| F201=*ICL1*-RM-R1 | TAATTATGTAGTCCAGCGCCTCTATCAAACTCGTTAAATATATTT TGCTAAACAAACTGCTGGGTTAGTACGGGACAGATGTTTGATGT GACACTTGCATTGGTGAGACCGACTTCTTTG | Repair fragment for deletion |
| F259=User-p*BR30*-*AtCAD*-PF | CGTGCGAUcgaactgtgtaggaatgaaac | Promoter amplification |
| F260=User-p*BR30*-*AtCAD*-PR | ATTGTGGCggUTGTGTGTGGGTGTGTCGTTCG | Promoter amplification |
| F261=User-p*BR30*-*AtCAD*-GF | accGCCACAAUGACCAAGCAGTCTGCCGACTC | Gene amplification |
| F262=User-*ICL1*-*AtCAD*-PF | CGTGCGAUgtagcgttggtctgtcctgtcg | Promoter amplification |
| F263=User-*ICL1*-*AtCAD*-PR | ATTGTGGCUTGGGTTAGTACGGGACAGATG | Promoter amplification |
| F264=User-*ICL1*-*AtCAD*-GF | aGCCACAAUGACCAAGCAGTCTGCCGACTC | Gene amplification |
| F322=U1-4UASTef-PF | CGTGCGAUCGATACGCGTatcgatacgc | Promoter amplification |
| F323=U1-4UASTef-PR | ATTGTGGCUgtggatccttcgggtgtgag | Promoter amplification |
| F324=U1-4UAS/Tef-*AtMTT*-GF | aGCCACAAUGGACAGCAAGATTCAGAC | Gene amplification |
| F325=U1-4UAS/Tef-*AtMTT*-GR | CACGCGAUTTAGTTGGGCTGTGTCAGGAAC | Gene amplification |
| F326=U1-4UAS/Tef-UmMtt1-GF | aGCCACAAUGCCTCCTAGCGGCCGAAAGG | Gene amplification |
| F327=U1-4UAS/Tef-UmMtt1-GR | CACGCGAUTCAGCTCTCGGGACCAGCGAGC | Gene amplification |
| F328=U1-4UAS/Tef-AkCtp1-GF | aGCCACAAUGGCTACTTCCGAAAACGACAAG | Gene amplification |
| F329=U1-4UAS/Tef-AkCtp1-GR | CACGCGAUTCAAATGTATCGTCGCTCGGGG | Gene amplification |
| F344=U1-Tefin-PR | AGTACUGCAAAAAGTGCTGGTCGGATG | Promoter amplification |
| F346=U1-Tefin-*AtCAD*-GF | AGTACUAACCGCAGACCAAGCAGTCTGCCGACTC | Gene amplification |
| F347=U1-Tefin-*AtMTT*-GF | AGTACUAACCGCAGGACAGCAAGATTCAGACTAAC | Gene amplification |
| F348=U1-Tefin-Ummtt1-GF | AGTACUAACCGCAGCCTCCTAGCGGCCGAAAGGTG | Gene amplification |
| F349=U1-Tefin-AkCtp1-GF | AGTACUAACCGCAGGCTACTTCCGAAAACGACAAG | Gene amplification |
| F350=U1-Tefin-*AtACO*d-GF | AGTACUAACCGCAGGCTACCGTGGCCGACTCGCCC | Gene amplification |
| F351=U1-Tef-cox4/CAD-GSG-ACO/d-1R | AGGGGAGACUTTACACCAGGGGAGACTTCACG(wrongreorder)-> | *AtACO* and *AtCAD* fused protein |
| F352=U1-Tef-cox4/CAD-GSG-ACO/d-2F | AGTCTCCCCUGGTGGGTTCTGGTGCTACCGTGGC CGACTCGCC | *AtACO* and *AtCAD* fused protein |

| | | |
|---|---|---|
| F353=U1-Tef-ACO/d-GSG-CAD-1R | AGAACCGTUGGAGGCGGCCTTTCGAGCCATG | *AtACO* and *AtCAD* fused protein |
| F354=U1-Tef-ACO/d-GSG-CAD-2R | AACGGTTCUGGTATGACCAAGCAGTCTGCCGA | *AtACO* and *AtCAD* fused protein |
| F367=U1-4UAS/Tef-SceDIC1-GF | aGCCACAAUGAGCACAAATGCCAAGGAAAG | Gene overexpression |
| F368=U1-4UAS/Tef-SceDIC1-GR | CACGCGAUTCACTTGTCCTCCTTAGGCATG | Gene overexpression |
| F369=U1-4UAS/Tef-MmuSlc-GF | aGCCACAAUGGCTGAAGCCCGCACGTCGCG | Gene overexpression |
| F370=U1-4UAS/Tef-MmuSlc-GR | CACGCGAUTCAGGTCGTGGGAACCTTGATG | Gene overexpression |
| F371=U1-4UAS/Tef-AtDIC-GF | aGCCACAAUGGCTGAAGAGAAGAAAGCTC | Gene overexpression |
| F372=U1-4UAS/Tef-AtDIC-GR | CACGCGAUTCACATTCCAATCTTCTTCTG | Gene overexpression |
| F373=U1-Tefin-SceDIC1-GF | AGTACUAACCGCAGAGCACAAATGCCAAGGAAAGC | Gene overexpression |
| F374=U1-Tefin-MmuSlc-GF | AGTACUAACCGCAGGCTGAAGCCCGCACGTCGCG | Gene overexpression |
| F375=U1-Tefin-AthDIC-GF | AGTACUAACCGCAGGCTGAAGAGAAGAAAGCTCC | Gene overexpression |
| F400=pICD1-UF | ttgctctgcagaagaagaac | *IDH1* deletion |
| F401=pICD1-UR | cggctcggttttatatccgg | *IDH1* deletion |
| F402=pICD1-LF | atgctcaaccttagaaccgc | *IDH1* deletion |
| F403=pICD1-LR | ttgaaggttcgtcgcagagtc | *IDH1* deletion |
| F404=pICD1-FsnF | gagtctatggctgggatctc | *IDH1* deletion |
| F405=pICD1-FsnR | gacccttgagggcaaccttg | *IDH1* deletion |
| F426=ICD1-check-FF | tttagggaaaagccggtgggg | *IDH1* deletion |
| F427=ICD1-check-RR | gaccctgtactttgacagctc | *IDH1* deletion |
| F428=ICD2-check-FF | ctgttctcggttgttcaattg | *IDH1* deletion |
| F429=ICD2-check-RR | tcaggatcacgattttgatag | *IDH1* deletion |
| F430=4UASTef-seam-PF | CACGCGAUCGATACGCGTatcgatacgcg | 4UASTef promoter |
| F431=4UASTef-seam-PF | ACCTGCACUtgtggatccttcgggtgtgag | 4UASTef promoter |
| F432=4UASTef-seam-*AtMTT*-GF | AGTGCAGGUGCCACAATGGACAGCAAGATTCAGAC | *AtMTT* under 4UASTef promoter |
| F433=4UASTef-seam-*AtMTT*-GR | CGTGCGAUTTAGTTGGGCTGTGTCAGGAAC | *AtMTT* under 4UASTef promoter |
| F436=pICD1-1A00683-ck1 | TCTCAGAATCGCAATTTGCAGA | Check primer for*IDH1* promoter change |
| F437=pICD1-1D13716-ck2 | gctacttgtagtgggagaggg | Check primer for*IDH1* promoter change |
| F438=pICD1-1B21134-ck3 | ggtatgtatctctcctacccgg | Check primer for*IDH1* promoter change |
| F439=pICD1-1F27529-ck4 | cccgaagcgctgacatacttca | Check primer for*IDH1* promoter change |
| F440=pICD1-1E16597-ck5 | gatggactcccaggtgtacac | Check primer for*IDH1* promoter change |
| F441=pICD1-1B10130-ck6 | cctattctgcgagtgtcgtgg | Check primer for*IDH1* promoter change |
| F442=pICD1-1D07348-ck7 | catgcccacaacatgaacaag | Check primer for*IDH1* promoter change |
| F443=pICD1-1C24124-ck8 | ggcttgttccttgcaaccgag | Check primer for*IDH1* promoter change |

| | | |
|---|---|---|
| F444=pICD1-1F04643-ck9 | tgtctagtcacgtgtaggtgg | Check primer for*IDH1* promoter change |
| F445=pICD1-1A14199-ck10 | ggttcttcgacaaagccacac | Check primer for*IDH1* promoter change |
| F446=pICD1-1D13716-ampF | gtggtcattgaggtagacattg | Check primer for*IDH1* promoter change |
| F447=pICD1-1D13716-ampR | cgtttacgatcttgagcgaagc | Check primer for*IDH1* promoter change |
| F509=U-pTef(20bp)-F | CGTGCGAUAGAGACCGGGTTGGCGGCGC | Tef promoter |
| F510=U-pTef(20bp)-R | ATTTTGAAUGATTCTTATACTCAGAAGG | Tef promoter |
| F535=U-UmAdi1-F | ATTCAAAAUGCTCCACCCCATCGATACTAC | Gene overexpression |
| F536=U-UmAdi1-R | CACGCGAUTCACGACAAGCTTCGGTCAG | Gene overexpression |
| F537=U-UmTad1-F | ATTCAAAAUGGCTCCCGCCCTTAACGCCAAC | Gene overexpression |
| F538=U-UmTad1-R | CACGCGAUTCAGGCAGAAGACGGGCGGCTAAG | Gene overexpression |
| F546=4UASTef-noseam-PF | CGTGCGAUCGATACGCGTatcgatacgcg | 4UASTef promoter |
| F547=4UASTef-noseam-PR | ATtgtggaUccttcgggtgtgagttgac | 4UASTef promoter |
| F548=4UASTefnosm*AtMTT*-GF | atccacaAUGGACAGCAAGATTCAGACTAAC | *AtMTT* under 4UASTef promoter |
| F549=4UASTefnosm*AtMTT*-GR | CACGCGAUTTAGTTGGGCTGTGTCAGGAAC | *AtMTT* under 4UASTef promoter |
| F550=4UASTefnosm*AtCAD*-GF | atccacaAUGACCAAGCAGTCTGCCGACTC | *AtCAD* under 4UASTef promoter |
| F551=4UASTefnosm*AtCAD*-GR | CACGCGAUTTACACCAGGGGAGACTTCAC | *AtCAD* under 4UASTef promoter |
| F552=Ri-*IDH1*F1U0 | CGTGCGAUAGAGACCGGGTTGGCGGCGC | RNAi repeat |
| F553=Ri-*IDH1*F1D1 | AAGCATGGUCACGGGAAAGGAGCCCTCCACCCTCACGTtcccgctccttctccgagtcg | RNAi repeat |
| F554=Ri-*IDH1*F1D2 | AAGCATGGUCACGGGAAAGGAGCCCTCCACCCTCACGTccaattattccgctgcgtctc | RNAi repeat |
| F555=Ri-*IDH1*F1D3 | AAGCATGGUCACGGGAAAGGAGCCCTCCACCCTCACGTttgaaggttcgtcgcagagtc | RNAi repeat |
| F556=Ri-*IDH1*F1D4 | AAGCATGGUCACGGGAAAGGAGCCCTCCACCCTCACGTcttgagtcgcttgataatctg | RNAi repeat |
| F557=Ri-*IDH1*F2U1 | ACCATGCTUCCGGGAGACGGTGTGGGGCCTGAGCTGATGtcccgctccttctccgagtc | RNAi repeat |
| F558=Ri-*IDH1*F2U2 | ACCATGCTUCCGGGAGACGGTGTGGGGCCTGAGCTGATGccaattattccgctgcgtctc | RNAi repeat |
| F559=Ri-*IDH1*F2U3 | ACCATGCTUCCGGGAGACGGTGTGGGGCCTGAGCTGATGttgaaggttcgtcgcagagtc | RNAi repeat |
| F560=Ri-*IDH1*F2U4 | ACCATGCTUCCGGGAGACGGTGTGGGGCCTGAGCTGATGcttgagtcgcttgataatctg | RNAi repeat |
| F561=Ri-*IDH1*F2D0 | CACGCGAUatgctcaaccttagaaccgc | RNAi repeat |
| F562=Ri-*IDH1*F1D0 | ATTTGAATGAUTCTTATACTCAGAAGGAAATG | RNAi repeat |
| F563=Ri-*IDH1*F2U5 | ATCATTCAAAUttcttgtcgccaacttcgtg | RNAi repeat |
| F564=Ri-*IDH1*F2D5 | AAGCATGGUCACGGGAAAGGAGCCCTCCACCCTCACGTcgcaatccgcatttaaaaag | RNAi repeat |
| F565=Ri-*IDH1*F3U5 | ACCATGCTUCCGGGAGACGGTGTGGGGCCTGAGCTGATGcgcaatccgcatttaaaaag | RNAi repeat |
| F566=Ri-*IDH1*F3D5 | CACGCGAUtttcttgtcgccaacttcgtg | RNAi repeat |
| F567=Ri-*IDH1*F2U6 | ATCATTCAAAUcaaccttagaaccgcccttc | RNAi repeat |
| F568=Ri-*IDH1*F2D6 | AAGCATGGUCACGGGAAAGGAGCCCTCCACCCTCACGTattgtcccgctccttctccg | RNAi repeat |
| F569=Ri-*IDH1*F3U6 | ACCATGCTUCCGGGAGACGGTGTGGGGCCTGAGCTGATGattgtcccgctccttctccg | RNAi repeat |
| F570=Ri-*IDH1*F3D6 | CACGCGAUtcaaccttagaaccgcccttc | RNAi repeat |
| F571=Ri-*IDH1*F2U7 | ATCATTCAAAUgagtatctcggagcccggg | RNAi repeat |
| F572=Ri-*IDH1*F2D7 | AAGCATGGUCACGGGAAAGGAGCCCTCCACCCTCACGTggtctctcatcaccctcagttc | RNAi repeat |

| | | |
|---|---|---|
| F573=Ri-*IDH1*F3U7 | ACCATGCTUCCGGGAGACGGTGTGGGGCCTGAGCTGATggtctctcat caccctcagttc | RNAi repeat |
| F574=Ri-*IDH1*F3D7 | CACGCGAUtgagtatctcggagcccggg | RNAi repeat |
| F575=Ri-*IDH1*F2U8 | ATCATTCAAAUcgcaatccgcatttaaaaag | RNAi repeat |
| F576=Ri-*IDH1*F2D8 | CACGCGAUtttcttgtcgccaacttcgtg | RNAi repeat |
| F577=Ri-*IDH1*F2U9 | ATCATTCAAAUctacttgagtcgcttgataatc | RNAi repeat |
| F578=Ri-*IDH1*F2D9 | CACGCGAUatgctcaaccttagaaccgc | RNAi repeat |
| F579=Ri-*IDH1*F2U10 | ATCATTCAAAUgaaggttcgtcgcagagtcag | RNAi repeat |
| F580=Ri-*IDH1*F2D10 | CACGCGAUtcgactgggaggttgtcgacg | RNAi repeat |
| F613=P1F-3*C&M | CGTGCGAUTTACACCAGGGGAGACTTCACG | JFEC127 and JFEC159 |
| F614=P1R-3*C&M | ACCTGACACUCAGATGCATTCTTGGGCGGTC | JFEC127 and JFEC159 |
| F615=P2F-3*C&M | AGTGTCAGGUACGCAACTAACATGAATGAATACG | JFEC127 and JFEC159 |
| F616=P2R-3*C&M | ACAGGACTGUCAGATGCATTCTTGGGCGGTC | JFEC127 and JFEC159 |
| F617=P3F-3*C&M | ACAGTCCTGUACGCAACTAACATGAATGAATACG | JFEC127 and JFEC159 |
| F618=P3R-3*C&M | CACGCGAUTTAGTTGGGCTGTGTCAGGAAC | JFEC127 and JFEC159 |
| F637=ampd-UF | CGTGCGAUTATACACAAGAACCAGCATG | Ampd amplification |
| F638=ampd-UR | AGTCGAGGCUATTGTATATGAGTATCAAGCTC | Ampd amplification |
| F639=AMPDTEF-PF | AGCCTCGACUAGAGACCGGGTTGGCGGCGC | Ampd amplification |
| F640=AMPDTEF-PR | ATTGCTTGCTGCGGCAUTTTGAATGATTCTTATACTC | Ampd amplification |
| F641=ampd-DF | ATGCCGCAGCAAGCAAUGGATATCAAGGGCAAGGCC | Ampd amplification |
| F642=ampd-DR | CACGCGAUGATAGACACAGGTAGGATCTC | Ampd amplification |
| F669=JFEC127-c1ckR | TTAGTTGCGTACCTGACACTC | Check primer for JFEC127 and JFEC159 |
| F670=JFEC127-c2ckF | ATGCATCTGAGTGTCAGGTAC | Check primer for JFEC127 and JFEC159 |
| F671=JFEC127-c2ckR | GTTAGTTGCGTACAGGACTGT | Check primer for JFEC127 and JFEC159 |
| F672=JFEC127-c3ckF | GAATGCATCTGACAGTCCTGT | Check primer for JFEC127 and JFEC159 |
| F729=AMPD-cr3F | AAGAGGGGCTAATAACCTGGgttttagagct | gRNA expression cassette |
| F730=AMPD-cr3R | CCAGGTTATTAGCCCCTCTTtaaccaacct | gRNA expression cassette |
| F731=AMPD-cr4F | TTGGAGATAGAGCCGTTGGAgttttagagct | gRNA expression cassette |
| F732=AMPD-cr4R | TCCAACGGCTCTATCTCCAAtaaccaacct | gRNA expression cassette |

Table S4 IA production in different microorganisms

| Strain | Strategy for strain engineering | Carbon source | Fermentation type and tips | Titer (g/L) | Yield (mol/mol) | Productivity (g/L/h) | Produced IA amount (g) | Ref |
|---|---|---|---|---|---|---|---|---|
| ***A. terreus*** **DSM 23081** | Wild type | Glucose | Fed-batch in 1.5 L bioreactor, pH at 3.4 in production phase | 160 | 0.46 | 0.99 | NA | [4] |
| ***U. maydis*** **MB215** | Overexpression of native rai1 and *mttA* from *A. tereus*; deletion of cyp3 and fuz7 coding itaconate oxidase | Glucose | Fed-batch in 5 L bioreactor; in situ product crystallization with $CaCO_3$. | 220 | 0.33 | 0.46 | NA | [5] |
| ***A. niger*** **AB 1.13** | Overexpression of *acl*2, *citB*, *cadA*, *mttA* and *mfsA* | Glucose | Fed-batch in 10L bioreactor | 42.7 | 0.26 | 0.18 | NA | [6] |
| ***Saccharomyces cerevisiae*** **XYY286** | Overexpression of *cadA* from *A. tereus* and *acc2*; Deletion of *hxk2* | Glucose | Flask | 0.535 | - | - | NA | [7] |
| ***E. coli*** **ita36A** | Overexpression of *cadA* from *A. tereus*; native *icd* under temperature control; deletion of *aceA*, *pta*, *pykF pykA* | Glucose | Fed-batch in 1L bioreactor; temperature controlled two-stage process | 47 | 0.62 | 0.39 | NA | [8] |
| ***Corneybacterium glutamicum*** | Heterologous expression of *cadA*, from *A. terreus*; mutated icd variant for reduced isocitrate dehydrogenase activity | Acetate | Fed-batch in 42L bioreactor; pH and DO-coupled feeding strategy. | 29.2 | | 0.63 | NA | [9] |
| ***Yarrowia lipolytica*** | Cytosolic coexpression of CAD and ACOnoMLS | Glucose | Fed-batch | 4.6 | 0.058 | 0.027 | NA | [10] |
| ***Yarrowia lipolytica*** | Heterologous expression of *cadA*, *acoA*, *mttA* and *mfsA* from *A. terreus* | Glucose | Fed-batch | 22 | 0.056 | 0.05 | NA | [11] |
| ***Yarrowia lipolytica*** | Heterologous expression of CAD-ePTS1 and POT1, and deletion of ICL | Cooking oil | Batch in 5L bioreactor | 54.55 | NA | 0.568 | NA | [12] |
| ***Yarrowia lipolytica*** **JFYL122** | | Glucose | Fed-batch in 1 L bioreactor | 130.2 | 0.320 | 0.226 | 103.3 | This study |
| ***Yarrowia lipolytica*** **JFYL122** | | Glucose | Fed-batch in 50 L bioreactor | 94.1 | 0.360 | 0.246 | 6161.88 | This study |

Table S5 Media with varying Carbon / Nitrogen (C/N) ratios, PL, and SL conditions

| Media | Comments | Note |
|---|---|---|
| 10 g/L $(NH_4)_2SO_4$, 3 g/L $KH_2PO_4$, 0.5 g/L $MgSO_4$•$7H_2O$, 2 mL trace metals solution stock, and 1 mL of vitamin solution stock, 100 g/L initial glucose. 650 g/L glucose was fed when residual glucose is below 20 g/L. | C/N=22 | Fig. 6a |
| 10 g/L $(NH_4)_2SO_4$, 19.1 g/L $KH_2PO_4$, 10.4 g/L $K_2HPO_4$, 0.5 g/L $MgSO_4$•$7H_2O$, 39 g/L MES (0.2M), 2 mL trace metals solution stock, and 1 mL of vitamin solution stock | NR | Fig. 6b |
| 2.5 g/L $(NH_4)_2SO_4$, 19.1 g/L $KH_2PO_4$, 10.4 g/L $K_2HPO_4$, 0.5 g/L $MgSO_4$•$7H_2O$, 39 g/L MES (0.2M), 2 mL trace metals solution stock, and 1 mL of vitamin solution stock | NL | Fig. 6b |
| 10 g/L $(NH_4)_2SO_4$, 19.1 g/L $KH_2PO_4$, 0.15 g/L $K_2HPO_4$, 0.5 g/L $MgSO_4$•$7H_2O$, 39 g/L MES (0.2M), 2 mL trace metals solution stock, and 1 mL of vitamin solution stock | PL | Fig. 6b |
| 8 g/L $(NH_4)_2Cl$, 19.1 g/L $KH_2PO_4$, 10.4 g/L $K_2HPO_4$, 0.005 g/L $MgSO_4$•$7H_2O$, 39 g/L MES (0.2M), 2 mL trace metals solution stock, and 1 mL of vitamin solution stock | SL | Fig. 6b |
| 10 g/L $(NH_4)_2SO_4$, 3 g/L $KH_2PO_4$, 0.5 g/L $MgSO_4$•$7H_2O$, 2 mL trace metals solution stock, and 1 mL of vitamin solution stock, 100 g/L initial glucose. 650 g/L glucose was fed when residual glucose is below 20 g/L. | NR->NL | Fig. 6c |
| 2.5 g/L $(NH_4)_2SO_4$, 3 g/L $KH_2PO_4$, 0.5 g/L $MgSO_4$•$7H_2O$, 2 mL trace metals solution stock, and 1 mL of vitamin solution stock, 100 g/L initial glucose. 650 g/L glucose was fed when residual glucose is below 20 g/L. | NL | Fig. 6c |
| 10 g/L $(NH_4)_2SO_4$, 0.2 g/L $KH_2PO_4$, 0.5 g/L $MgSO_4$•$7H_2O$, 2 mL trace metals solution stock, and 1 mL of vitamin solution stock, 100 g/L initial glucose. | PL | Fig. 6c |
| 10 g/L $(NH_4)_2SO_4$, 3 g/L $KH_2PO_4$, 0.1 g/L $MgSO_4$•$7H_2O$, 2 mL trace metals solution stock, and 1 mL of vitamin solution stock, 100 g/L initial glucose. | SL | Fig. 6c |
| 10 g/L $(NH_4)_2SO_4$, 3 g/L $KH_2PO_4$, 0.5 g/L $MgSO_4$•$7H_2O$, 2 mL trace metals solution stock, and 1 mL of vitamin solution stock, 100 g/L initial glucose. 650 g/L glucose was fed when residual glucose is below 20 g/L. 2.5, 1, 0.1, and 0 g/L yeast extract was added for different conditions. | Test the effect of yeast extract | Fig. 6d |
| 10 g/L $(NH_4)_2SO_4$, 3 g/L $KH_2PO_4$, 0.5 g/L $MgSO_4$•$7H_2O$, 2 mL trace metals solution stock, and 1 mL of vitamin solution stock, 100 g/L initial glucose. 650 g/L glucose with 30 g/L yeast extract was fed continuously. | pH effect test | Fig. 6e |
| 7.5 g/L $(NH_4)_2SO_4$, 19.1 g/L $KH_2PO_4$, 10.4 g/L $K_2HPO_4$, 0.5 g/L $MgSO_4$•$7H_2O$, 39 g/L MES (0.2M), 2 mL trace metals solution stock, and 1 mL of vitamin solution stock with various concentrations of IA (0, 5, 10, 20, 40 g/L) or 20 g/L NaCl as a negative control to test the IA tolerance. The initial pH was 6.5. | IA tolerance test | Fig. 6f |
| 10 g/L $(NH_4)_2SO_4$, 3 g/L $KH_2PO_4$, 0.5 g/L $MgSO_4$•$7H_2O$, 2 mL trace metals solution stock, and 1 mL of vitamin solution stock, 100 g/L initial glucose. 650 g/L glucose with 30 g/L yeast extract was fed when residual glucose is below 20 g/L. | NR->NL | Fig. 6g |
| 2.5 g/L $(NH_4)_2SO_4$, 3 g/L $KH_2PO_4$, 0.5 g/L $MgSO_4$•$7H_2O$, 2 mL trace metals solution stock, and 1 mL of vitamin solution stock, 100 g/L initial glucose. 650 g/L glucose with 30 g/L yeast extract was fed when residual glucose is below 20 g/L. | NL | Fig. 6g |
| 10 g/L $(NH_4)_2SO_4$, 0.2 g/L $KH_2PO_4$, 0.5 g/L $MgSO_4$•$7H_2O$, 2 mL trace metals solution stock, and 1 mL of vitamin solution stock, 100 g/L initial glucose. 650 g/L glucose with 30 g/L yeast extract was fed when residual glucose is below 20 g/L. | PL | Fig. 6g |
| 10 g/L $(NH_4)_2SO_4$, 3 g/L $KH_2PO_4$, 0.1 g/L $MgSO_4$•$7H_2O$, 2 mL trace metals solution stock, and 1 mL of vitamin solution stock, 100 g/L initial glucose. 650 g/L glucose with 30 g/L yeast extract was fed when residual glucose is below 20 g/L. | SL | Fig. 6g |
| 10 g/L $(NH_4)_2SO_4$, or 4.5 g/L urea, 3 g/L $KH_2PO_4$, 0.5 g/L $MgSO_4$•$7H_2O$, 2 mL trace metals solution stock, and 1 mL of vitamin solution stock, 100 g/L initial glucose. 650 g/L glucose with 30 g/L yeast extract was fed continuously. Nitrogen source was fed during fermentation. | | Fig. 6h |
| 9 g/L urea, 6 g/L $KH_2PO_4$, 1 g/L $MgSO_4$•$7H_2O$, 4 mL trace metals solution stock, and 2 mL of vitamin solution stock, 100 g/L initial glucose. 650 g/L glucose with 30 g/L yeast extract was fed continuously. 400 g/L urea was fed during fermentation. | | Fig. 6i |
